# Successful Modulation of Temporoparietal Junction Activity and Stimulus-Driven Attention by fNIRS-based Neurofeedback – a Randomized Controlled Proof-of-Concept Study

**DOI:** 10.1101/2023.03.13.532169

**Authors:** Simon H. Kohl, Pia Melies, Johannes Uttecht, Michael Lührs, Laura Bell, David M. A. Mehler, Surjo R. Soekadar, Shivakumar Viswanathan, Kerstin Konrad

## Abstract

The right temporoparietal junction (rTPJ) is a core hub in neural networks associated with reorienting of attention and social cognition. However, it remains unknown whether participants can learn to actively modulate their rTPJ activity via neurofeedback. Here, we explored the feasibility of functional near-infrared spectroscopy (fNIRS)-based neurofeedback in modulating rTPJ activity and its effect on rTPJ functions such as reorienting of attention and visual perspective taking. In a bidirectional regulation control group design, 50 healthy participants were either reinforced to up- or downregulate rTPJ activation over four days of training.

Both groups showed an increase in rTPJ activity right from the beginning of the trainingbut only the upregulation group maintained this effect, while the downregulation group showed a decline from the initial rTPJ activation. This suggests a learning effect in the downregulation exclusively, making it challenging to draw definitive conclusions about the effectiveness of rTPJ upregulation training. However, we observed group-specific effects on the behavioral level. We found a significant group x time interaction effect in the performance of the reorienting of attention task and group-specific changes, with decreased reaction times (RTs) in the upregulation group and increased RTs in the downregulation group across all conditions after the neurofeedback training. Those with low baseline performance showed greater improvements. In the perspective-taking task, however, only time effects were observed that were non-group-specific.These findings demonstrate that fNIRS-based neurofeedback is a feasible method to modulate rTPJ functions with preliminary evidence of neurophysiologically specific effects, thus paving the way for future applications of non-invasive rTPJ modulation in neuropsychiatric disorders.

**Graphical abstract:** 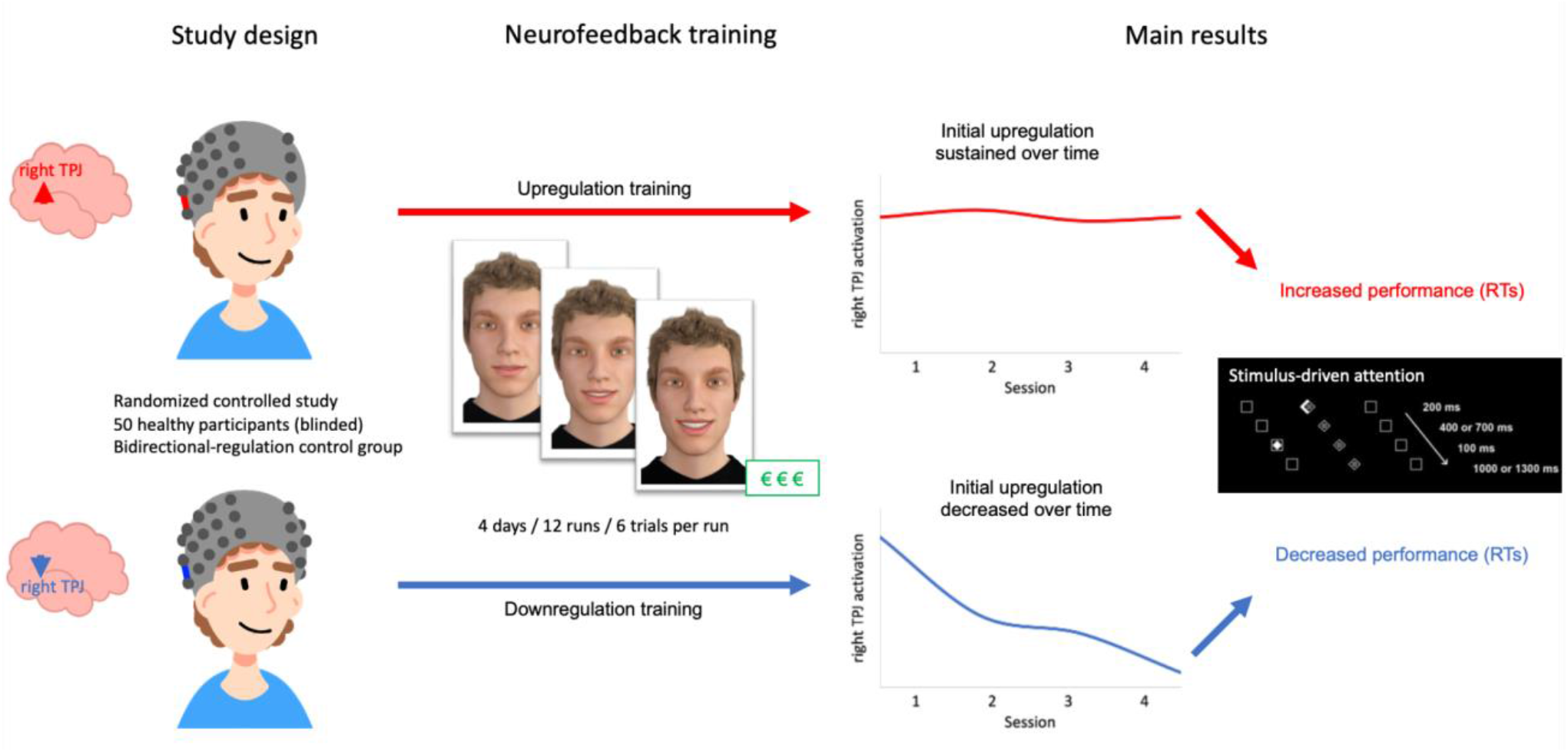

**Highlights:**

- the right temporoparietal junction (rTPJ) as a core hub for attentive and socio-cognitive functions is a promising target for neuromodulatory interventions
- first single-blinded, randomized controlled study demonstrates feasibility and effectiveness of the fNIRS-based neurofeedback training of the rTPJ in healthy adults
- subjects are able to regulate the rTPJ with different learning characteristics
- first evidence of a neurophysiologically specific effect on stimulus-driven attention
- findings have important implications for clinical translation of neurofeedback interventions targeting the rTPJ

## 1 Introduction

The right temporoparietal junction (TPJ) is considered a central hub of the human brain being involved in diverse mental functions. Theoretical models stress its involvement in stimulus-driven attention and social cognition and discuss its essential role in detecting violations of expectations, contextual updating, mental state shifting, and sense of agency (Decety and Lamm, 2007; Corbetta et al., 2008; van Overwalle, 2009; Geng and Vossel, 2013; Krall et al., 2015). Due to its diverse anatomical and functional connections, the TPJ is also considered an important brain region for communication with neighboring, partially overlapping networks, forming a potential hub where multiple networks converge and interact (Mars et al., 2012; Carter and Huettel, 2013).

Neuromodulation of such high degree network hubs or control points may result in greater changes in neural networks and associated behaviors and cognitive functions than neuromodulation of low degree nodes. Therefore, they are considered hot spots for targeted brain-based interventions (Murphy and Bassett, 2017)(Murphy and Bassett, 2017)(Murphy and Bassett, 2017).

Furthermore, targeting such high degree hubs using non-invasive neuromodulation, such as neurofeedback, is interesting from a translational perspective. Testing the causal role of the hub in the network by neuromodulation followed by observation of its behavioral/functional consequences may inform therapeutic interventions for brain disorders associated with this hub, e.g., autism spectrum disorder (ASD), depression and schizophrenia (Kana et al., 2015; Penner et al., 2018). In turn, testing these potential interventions will increase our understanding of this neural network hub and its role in the respective disorder.

Previous neuromodulation studies mostly relied on neurostimulation techniques such as transcranial magnetic stimulation (TMS) or transcranial direct current stimulation (tDCS) to disrupt or enhance TPJ functions while neurofeedback was utilized to a lesser extent.

TMS studies have demonstrated a decrease in spatial attention performance when disrupting activation in the right TPJ (rTPJ) (Krall et al., 2016; Mengotti et al., 2022). Conversely, a study conducted by Roy et al. (2015) used tDCS to enhance activation in the right posterior parietal cortex, which includes parts of the rTPJ, resulting in improved attention re-orienting following stimulation. Regarding socio-cognitive abilities such as visual perspective taking (vPT) and imitation control, the evidence for potential enhancement through tDCS is promising but mixed (Santiesteban et al., 2012, 2015; Nobusako et al., 2017; Yang et al., 2020). However, tDCS studies have reported no significant enhancing effects on other complex socio-cognitive abilities, including theory of mind (ToM; Santiesteban et al., 2015), empathy, emotion recognition, and joint attention (Pereira et al., 2021). In fact, inhibitory tDCS for ToM and empathy (Mai et al., 2016), as well as inhibitory TMS for ToM (Krall et al., 2016), have shown disruptive effects.

Together, these studies provide first evidence that neuromodulation of the rTPJ can be used to improve reorienting of attention and certain facets of socio-cognitive abilities, such as vPT. Therefore, the rTPJ may also be a promising target for neurofeedback interventions, offering potentially new treatment options for neuropsychiatric disorders characterized by deficient TPJ functions such as ASD (Esse Wilson et al., 2018; Salehinejad et al., 2021)

Neurofeedback based on functional near-infrared spectroscopy (fNIRS) is similar to neurostimulation a causative neuromodulation technique for modulating activation of circumscribed neocortical brain regions, although likely with less specific and more global effects on brain networks than neurostimulation. By providing real-time feedback of hemodynamic correlates of neural activity (e.g., changes in oxyhemoglobin), participants can learn to regulate the brain activity of specific target regions. In particular, fNIRS-based neurofeedback offers several advantages when it comes to clinical translation. It is an easy-to-use, non-invasive, and endogenous form of neuromodulation, which allows long-term learning through the reinforcement of neural activity and cognitive strategies with therapeutic potential. Moreover, it is safe and well tolerated, and is therefore associated with fewer ethical concerns than other neuromodulation techniques (Kohl et al., 2020; Soekadar et al., 2021). Across different studies, preliminary but compelling evidence suggests that the activation of a neural network, including the TPJ, can be successfully modulated by neurofeedback based on functional magnetic resonance imaging (fMRI) (Harmelech et al., 2015; Emmert et al., 2016; Direito et al., 2019, 2021; Pamplona et al., 2020. However, behavioral effects and specificity of findings are less clear, and no study has yet targeted rTPJ activity using fNIRS-based neurofeedback.

In the current study, we aimed to fill this gap and investigated the feasibility and effectiveness of fNIRS-based neurofeedback training employing social/monetary reward (Mathiak et al., 2015) to control rTPJ activity in healthy participants. We conducted a randomized, controlled proof-of-concept study employing a bidirectional-regulation control group design, which allows for the detection of neurophysiologically specific effects (Sorger et al., 2019). More specifically, we aimed to explore three aspects: (1) Can participants learn to increase/decrease the activity of the rTPJ using fNIRS-based neurofeedback and how is their learning behavior over the course of the training characterized? (2) Are there any specific behavioral effects in stimulus-driven attention and vPT following the neurofeedback training? (3) What are potential predictors of behavioral improvements? We hypothesized that healthy adult participants could gain control over activation of the rTPJ with fNIRS-based neurofeedback and that successful upregulation would be accompanied by improved performance in a reorienting of attention task and a vPT task. In contrast, we assumed that downregulation would be associated with either decreased performance or no change in performance. Based on previous findings on specific traits associated with rTPJ function, e.g., empathy and autistic traits (Kana et al., 2014; Donaldson et al., 2018; Yang et al., 2020), we tested predictors of behavioral change on an exploratory level. Due to the scarcity of neurofeedback studies targeting rTPJ activation, we identified a rather broad set of potential predictors without stating directed hypothesis for each of them (see methods section).

## 2 Methods

### 2.1 Participants

Fifty right-handed healthy participants (age 18-30 years) were recruited via flyer and social media announcements. Participants were screened during a telephone interview prior to participating in the study and were excluded if they had a history of psychiatric or neurological diseases, drug or alcohol abuse, or if they were undergoing current psychopharmacological or psychotherapeutic treatments. Participants were informed about the study procedure and signed an informed consent document. At the end of the study, they received a financial compensation of at least 60€ for attending all sessions, along with an additional monetary reward depending on the success of the neurofeedback training.

The study protocol was approved by the local ethics committee (EK 148/18) and conducted in accordance with the Declaration of Helsinki (World Medical Association, 2013)(World Medical Association, 2013), with the exception that it was not pre-registered on a publicly accessible database.

The participants were randomly allocated to the study groups, which were balanced out in terms of gender and the order of task assessments. Twenty subjects were allocated to the downregulation group and ten more (30 participants) to the upregulation group in order to provide higher statistical power for later subgroup analyses in this group.

After the first eleven participants, we noticed an error in the online preprocessing script (motion correction algorithm), which led to small deviations of the feedback displayed during the neurofeedback training. We simulated the feedback signal of these participants using the corrected script and calculated the accordance with the original feedback signal. Five participants (3 in the upregulation group, 2 in the downregulation group) showed an accordance below 90% and were therefore excluded from further analysis.

No a priori power analysis was conducted. However, according to a sensitivity analysis, a mixed analysis of variance (ANOVA) including 45 participants was sufficiently powered (80%) to detect a group x time interaction effect of at least *f* = 0.43 (assuming no violation of sphericity and a correlation among repeated measure of 0.8) or 0.77 for an independent *t*-test.

### 2.2 Study design

We applied a single-blinded, randomized controlled between-subject design. Participants were blinded to group assignments, but experimenters were not. We followed the recently published best practices for fNIRS publications (Yücel et al., 2021) and the consensus on the reporting and experimental design of clinical and cognitive-behavioral neurofeedback studies (CRED-nf checklist (Ros et al., 2020; see Supplementary Material 2)).

All participants took part in four appointments, including a pre- and post-assessment session with one additional short neurofeedback training session (day 1 and day 4) as well as two longer neurofeedback training sessions (day 2 and day 3; see Figure 1). The four appointments were scheduled within one week (*M* = 6.71 ± 2.23 days) and the pre- and post-assessment sessions at the same time of day. The procedure on each day was as follows:

**Figure 1.**
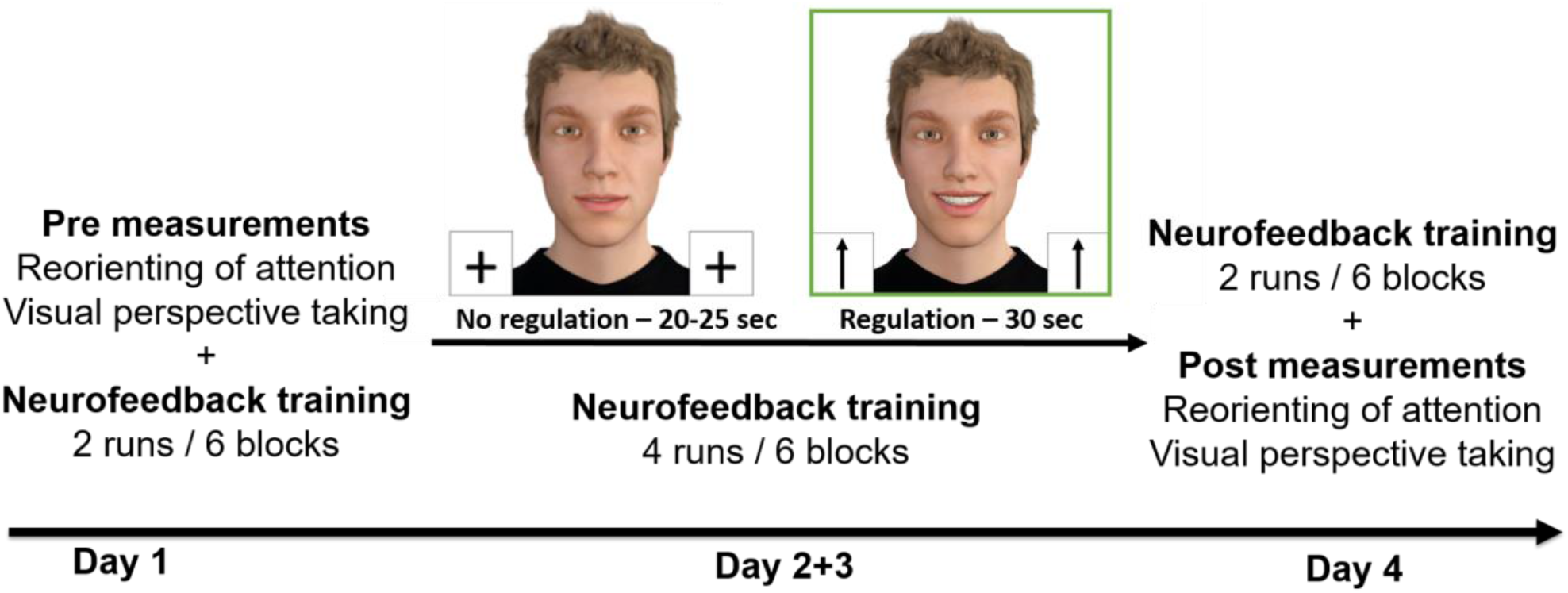
Study design Procedure

*Day 1:* To evaluate self-efficacy as a potential mechanism of neurofeedback effects and address group differences, participants completed the German version of the general self-efficacy scale (Schwarzer and Jerusalem, 1995). Additionally, a questionnaire was administered to assess the participants’ expectations and motivation towards the neurofeedback training, offering further controls for non-specific psychological mechanisms. After a short practice session, the pre-assessment of the (1) reorienting of attention task and (2) perspective-taking task took place. The order of these two tasks was counterbalanced across participants within both groups. Participants subsequently received specific instructions about the neurofeedback training and underwent two runs of neurofeedback training.

*Days 2 and 3:* On days 2 and 3, participants underwent neurofeedback sessions with four runs each. To assess pre-post changes in mood states and resting state brain activity, we assessed the German short version of the Profile of Mood States (McNair et al., 1981) and recorded a 10 min resting-state fNIRS measurement (to be reported elsewhere) before the training started on day 2 and after the training was completed on day 3. Between days 2 and 3, participants completed standardized questionnaires to account for variations in socio-cognitive traits among groups and predict neurofeedback effects. These traits, including autistic traits, empathy, cognitive styles, as well as ADHD symptoms have the potential to impact rTPJ functioning (Kana et al., 2014; Barman et al., 2015; Donaldson et al., 2018). All traits were assessed dimensionally using the German version of the Social Responsiveness Scale (Bölte, 2012), the Adult ADHD Self-Report Scale v1.1 (Kessler et al., 2005), the German version of the Interpersonal Reactivity Index (IRI; Davis, 1983), the autism-spectrum quotient (Baron-Cohen et al., 2001), the systemizing quotient (Baron-Cohen et al., 2003), and the empathy quotient (Baron-Cohen and Wheelwright, 2004).

*Day 4:* Participants underwent a short neurofeedback training session of two runs, followed by the post-assessment of the reorienting of attention and perspective-taking task. At the end of the session, participants filled in the general self-efficacy scale again as well as a debriefing questionnaire to further assess feasibility and unspecific mechanisms. This questionnaire included items assessing participants’ evaluation of the neurofeedback training, for example “I believe the training helped to improve my attention”, “I enjoyed the training”, “The experimenter was trustworthy”, etc. Furthermore, they were asked to guess the group condition they had been randomly assigned to. The fNIRS system was set up at the beginning of each day, except for day 1, which began with comprehensive instructions and practice runs. For all tasks, stimuli were presented on a 24-inch LCD screen (1920 x 1080 pixels) using the Psychtoolbox on Matlab 2017a (The Mathworks Inc, Natick, MA) and being run on a Windows PC. Participants viewed the screen at a distance of approximately 50cm. Responses were acquired using a standard keyboard.

#### Neurofeedback training

Participants were blinded to their group assignment and were told that, depending on their group assignments, the goal of the training was to increase or decrease activation of a specific brain region. Irrespective of group assignment, participants in both groups received the same instructions.

All participants received standardized information and instructions about the neurofeedback training (see Supplementary Material 1) based on Greer et al. (2014). They were instructed not to use any respiratory or motor strategies but to remain still, breathe regularly, and only rely on mental strategies to regulate their brain activity. The training took place on all four days and comprised 12 runs in total: two runs on days 1 and 4 (∼12 minute/day), and four runs on days 2 and 3 (∼25 minutes/day). Each neurofeedback run consisted of six blocks. Each block started with a 25s/30s no-regulation condition followed by a 30s regulation condition, and the block ended with a 2s reward presentation (see Figure 1). We varied the durations of the no-regulation condition to avoid synchronization with physiological confounds, such as breathing patterns and Mayer waves, during the task and to increase design efficiency (Kinoshita et al., 2016; Yücel et al., 2021). On each block, the face of a human avatar was continuously displayed on the screen. We used DAZ Studio 4.9 (DAZ Productions, Inc., USA) to create a modified version of the stimuli validated by Hartz et al. (2021). Eleven pictures of the avatar with different levels of smiling were created for the visualization of the feedback signal.

During the no-regulation condition, participants were instructed to passively look at the avatar, which maintained a neutral facial expression. During the regulation condition, real-time feedback of rTPJ activity was presented visually on a screen using a smiling avatar (social reward). Participants were instructed to regulate and make the avatar smile, which was modulated in real time by their rTPJ activation.

To foster motivation, participants received a monetary reward for successful regulation. In each regulation trial, the participant received 0.01€ per second exceeding a certain individual threshold (see real-time fNIRS data processing (online analysis)). Whenever participants exceeded this reward threshold, a green frame appeared around the feedback display, indicating that their regulation was earning an incentive. The total amount earned on each trial was presented on the screen at the end of the trial. This threshold was adapted according to individual regulation performance (see 0. for a detailed description).

In neurofeedback training, providing explicit mental strategies is not necessary but initially seems to facilitate learning (Scharnowski et al., 2015). Therefore, we provided some example strategies that could be helpful to regulate rTPJ activity (e.g., strategies related to ToM, empathy, thinking, imagination of positive events, counting, etc.; see Supplementary Material 1). However, participants were encouraged to find their own individual successful strategy by trial and error. After each neurofeedback run, we asked participants to verbally report which strategies they used and how successful they rated this strategy (Likert scale ranging from 1 to 5). After each session, we also assessed participants’ motivation to continue participating in the training and their beliefs about being able to control their brain activity.

#### Reorienting of attention task

Reorienting of attention is defined as the capacity to alter the focus of attention to unexpected, external stimuli while expecting another task/situation. We assessed the reorienting of attention using a modified version of the Posner paradigm (Posner, 1980; Vossel et al., 2009; Krall et al., 2016). In this task (see Figure 2), a central diamond (fixation point) was displayed between two horizontally arranged boxes. For each trial, a central cue was presented for 200ms indicating whether a target would appear on the right or the left side of the screen (brightening of the diamond to the right or left, respectively). After a variable cue-target interval of 400ms or 700ms, the target (white diamond) appeared for 100ms with a certain probability at the cued (valid cueing) or at the non-cued location (invalid cueing) and the participant had to indicate on which side it appeared by pressing a button using his/her right hand. The target-cue stimulus onset asynchrony was either 1000ms or 1300ms. All stimuli were presented on a black background. Since fNIRS was assessed during the task, the trials were presented in a blocked design. The task consisted of a total of twelve blocks, with six invalid blocks and six valid blocks. Each invalid block comprised twelve valid and eight invalid trials and each valid block comprised 20 valid trials only. Hence, the overall distribution of valid trials (192 of 240) and invalid trials (48 of 240) was 80% vs. 20%. The blocks were presented in a randomized order to mitigate anticipatory effects. Participants were told that the cue was not always informative, but they were not informed about the different blocks beforehand. The task blocks with a 40s duration were separated by 20s or 25s rest periods in which the same visual stimuli, but no cues or targets, were presented.

**Figure 2.**
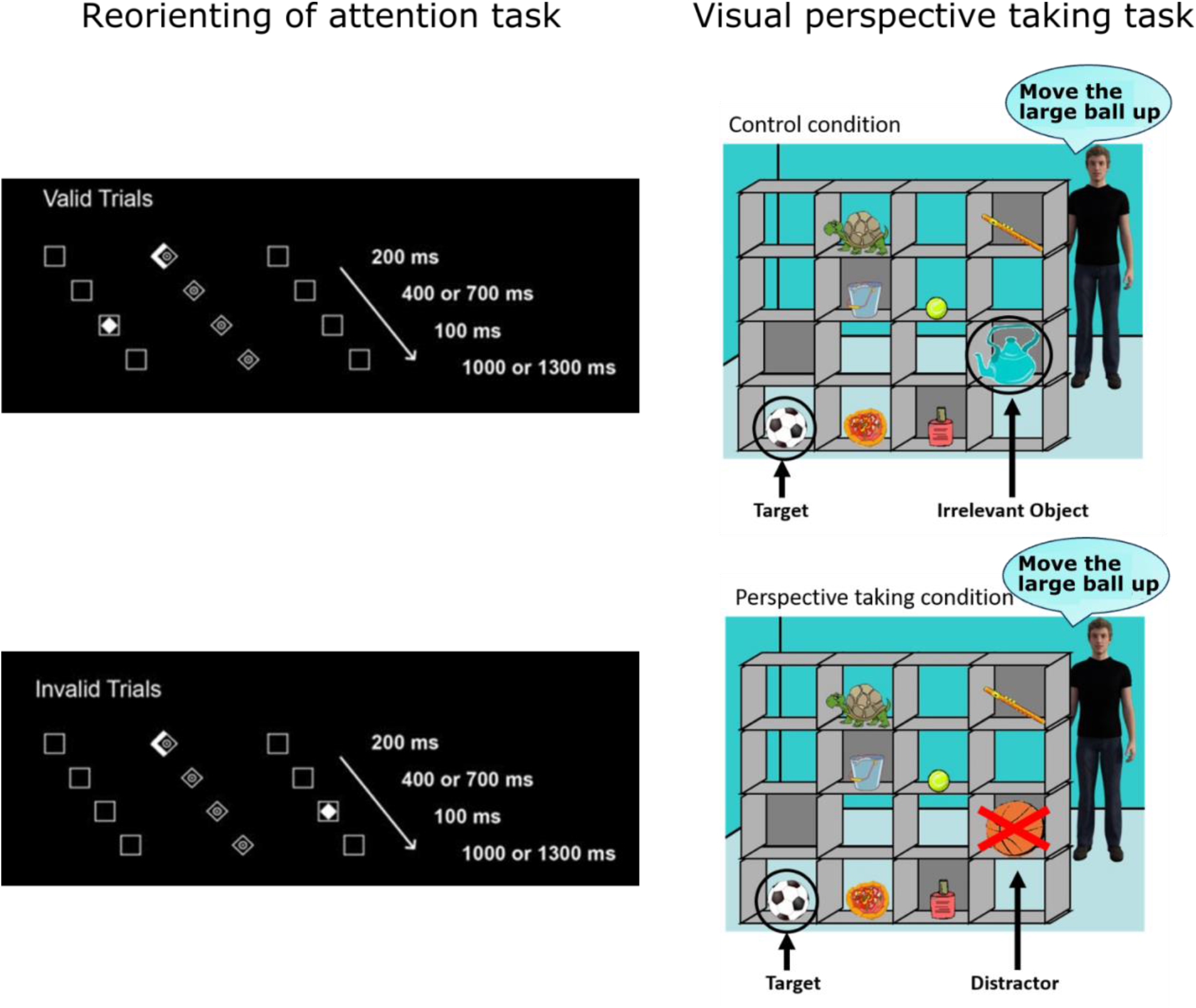
Illustration of experimental tasks used for pre-post measurements. The reorienting of attention task (left; adopted from Krall et al., 2016) and the visual perspective-taking task (right; adopted from Symeonidou et al. (2016) were used to measure the effects of the neurofeedback training. [*Note that in the perspective-taking task (right), the speech bubbles are only shown for illustration. Auditory instructions were provided to the participants by the director (see text).]*

#### Visual perspective-taking (vPT) task

vPT refers to the ability to infer spatial relationships between objects from different viewing angles. We assessed vPT with the widely used Director paradigm according to Dumontheil et al. (2010) and Symeonidou et al. (2016), as this task has been successfully used to assess the effects of tDCS stimulation of the TPJ (Santiesteban et al., 2012, 2015). In this task, participants saw a visual scene with a 4 x 4 set of shelves containing eight different objects (see Figure 2) and were instructed to take the perspective of a “director” standing behind the shelves and giving them auditory instructions to move certain objects on the shelves by clicking a mouse on the respective target object. Importantly, some of the objects were occluded from the view of the director, which participants had to take into account in order to respond correctly in the perspective taking (PT) condition. This can be seen in Figure 2 where the “director” refers to the football instead of the large basketball (distractor), which is occluded from his view. In the control condition (non-perspective-taking (NPT) condition), the distractor is replaced by an irrelevant object. For a more detailed description of this task, see Dumontheil et al. (2010) and Symeonidou et al. (2016). Each block consisted of four trials. The PT and NPT blocks were presented in a pseudo-randomized order in such a way that no more than two blocks of the same condition were presented consecutively. The task blocks (24s) were separated by a rest period with a duration of 20s or 25s. RTs were recorded from the onset of the auditory instruction to the participant’s mouse click response.

### 2.3 fNIRS acquisition

We used the ETG-4000 continuous wave system (Hitachi Medical Corporation, Tokyo, Japan) to measure changes in oxy-(HbO) and deoxyhemoglobin (HbR) concentrations at a rate of 10Hz with two wavelengths (695nm and 830nm). Two 3 × 5 probe sets (2 × 22 measurement channels) were placed bitemporally on the participant’s head to cover temporal and frontal brain regions and were attached using electroencephalography (EEG) caps (Easycap GmbH, Herrsching, Germany). The interoptode distance was 3cm. The probe sets were placed on the participants’ heads in such a way that the second most posterior optode of the lowest row was placed over T3/T4 of the EEG 10-20 system (Jasper, 1958)(Jasper, 1958) and the most anterior optode of the lowest row was placed approximately over the eyebrow (see Figure 3). If necessary, hair was moved away from optode holders in order to increase the quality of the signal. Furthermore, we instructed participants each day to stay relaxed, breathe regularly, and keep the movement of their heads to a minimum.

**Figure 3.**
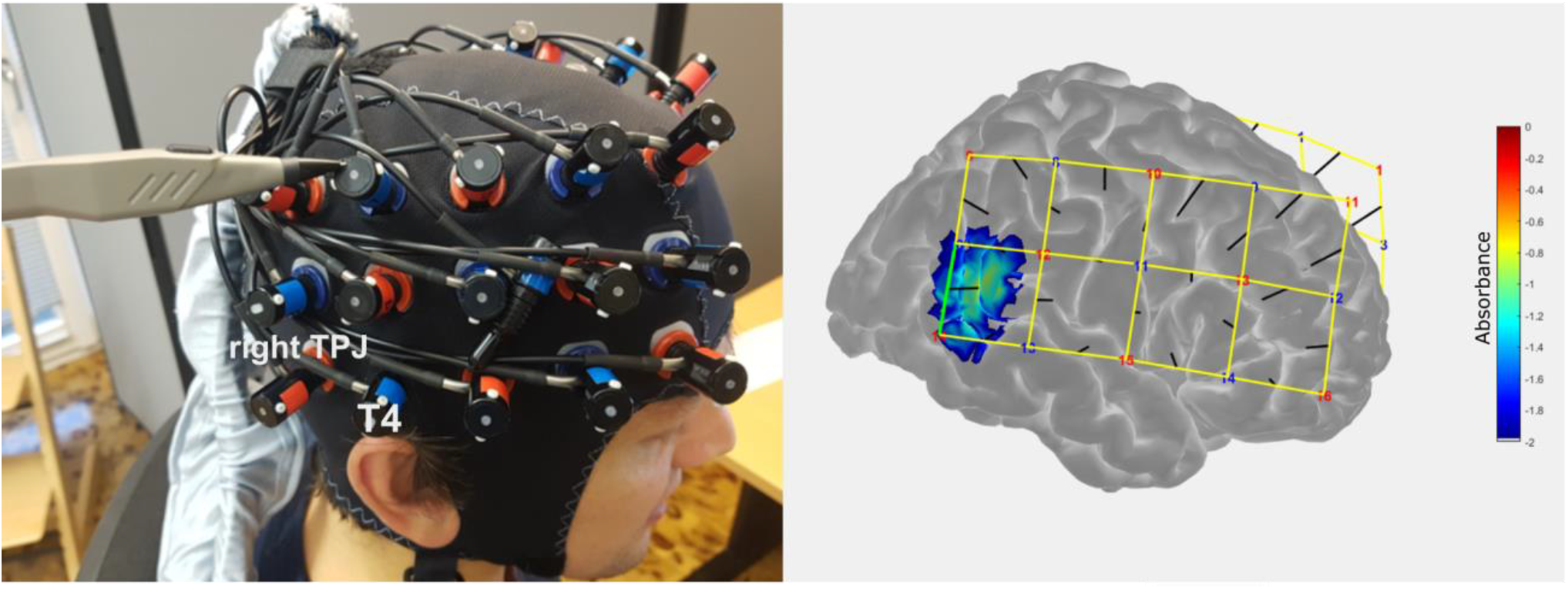
fNIRS optode arrangement and sensitivity profile for the feedback channel. The feedback channel corresponds to anterior parts of the rTPJ (MNI: x = 56 ± 6.4, y = −49 ± 4.6, z = 18 ± 6.9).

To select the best channel for the feedback processing that covers the rTPJ, prior to the current study, we conducted digitizer measurements using a Patriot 3D Digitizer (Polhemus, Colchester, Vermont) in a separate sample of five pilot participants wearing an fNIRS optode arrangement available from a previous study. In all five subjects, the same channel corresponded to anterior parts of the rTPJ (see Figure 3). To confirm the anatomical specificity of this channel, we also conducted digitizer measurements in all participants of the current study after each experimental session. Anatomical locations of the optodes in relation to standard head landmarks (nasion, inion, Cz, and preauricular points) were assessed. Cortical sensitivities of all channels were estimated through Monte Carlo photon migration simulations (1,000,000 photons) using AtlasViewer implemented in Homer v2.8 (Huppert et al., 2009; Aasted et al., 2015). Montreal Neurological Institute (MNI) coordinates for each subject and session were extracted and averaged for each participant. In total, 5% of data (i.e., 10 of the 200 samples obtained from 50 participants and 4 sessions) were excluded from this analysis due to errors during the digitizer measurements, which resulted in implausible estimations of MNI coordinates. The average MNI coordinate of the feedback channel (x = 56 ± 6.4, y = −49 ± 4.6, z = 18 ± 6.9) corresponded to anterior parts of the rTPJ, previously reported in a meta-analysis for reorienting of attention and theory of mind contrasts (Krall et al., 2015).

### 2.4 Real-time fNIRS data processing (online analysis)

Participants received feedback in real time about the instantaneous HbO activity at one single channel placed over the rTPJ (see section on fNIRS acquisition). The procedure to convert the HbO activity into feedback (updated every 100ms) involved several steps as described below (see Figure 4).

**Figure 4.**
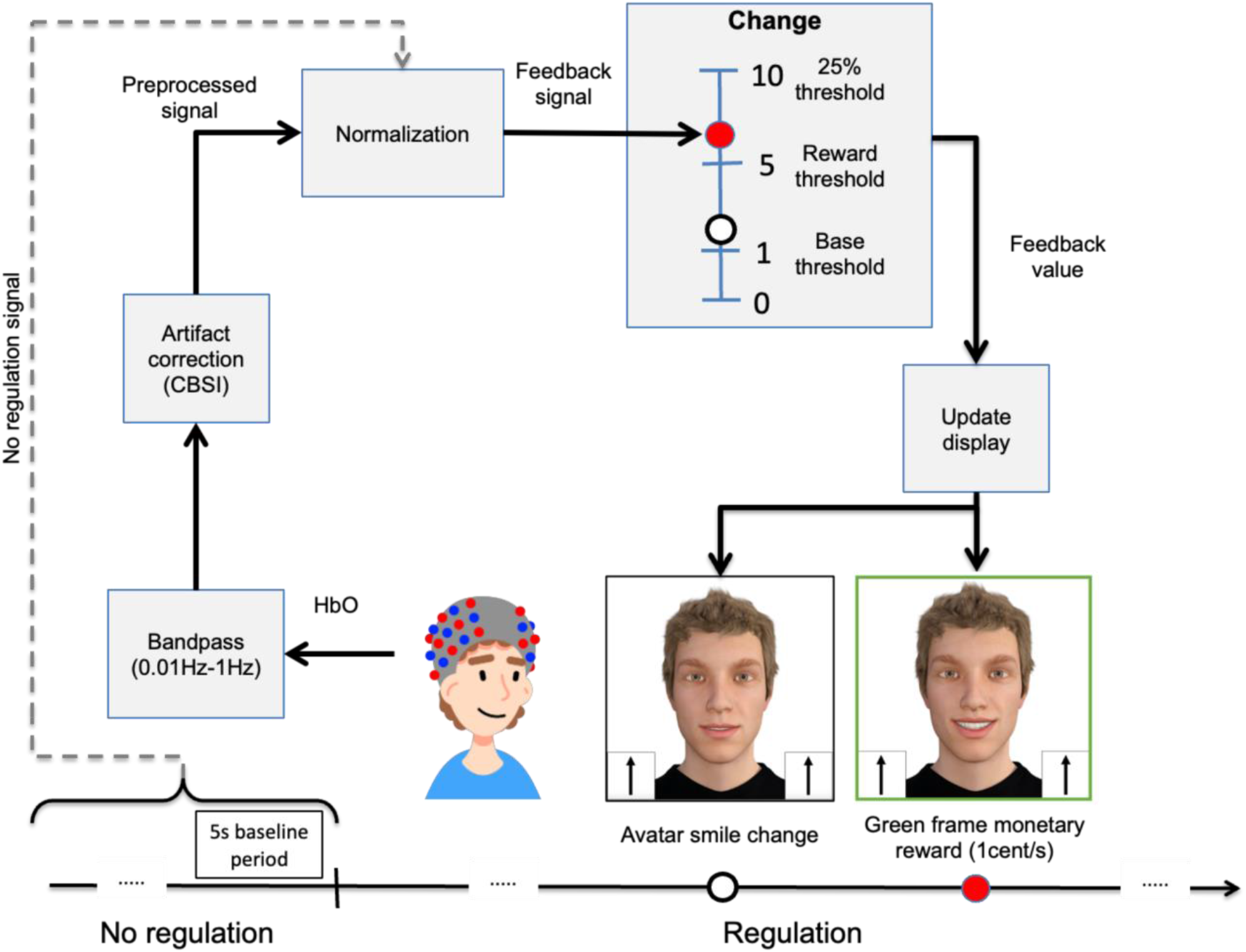
Real-time fNIRS data processing. Correlation-based signal improvement (CBSI) algorithm (Cui et al., 2010)(Cui et al., 2010).

The raw signal was first preprocessed by the ETG-4000 using a high-pass filter of 0.01Hz and a low-pass filter of 1Hz and a moving average of 5s. This preprocessed HbO signal was then sent in real time to an external computer where it was further processed using a customized Matlab script.

Motion artifacts in the signal were then removed using the correlation-based signal improvement (CBSI) algorithm (Cui et al., 2010). This algorithm calculates the corrected signal as a linear combination of HbO and HbR scaled by their standard deviations, based on the assumption that HbO and HbR are highly negatively correlated. Furthermore, the algorithm assumes that the signal has been offset-corrected to have zero mean.

Using the CBSI algorithm, the corrected HbO signal at time *t* (denoted as *X*_corr_ (t)) was obtained using the following expression:

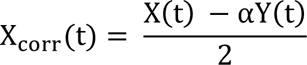

where *X*(*t*) and *Y*(*t*) are the measured values of the HbO and HbR values, respectively, at time *t* (after offset correction), and α is the ratio of the noise amplitude in the HbO and HbR signals. To estimate the noise amplitude ratio α and perform the offset correction, we used the HbO and HbR signals from the last 30s of the no-regulation period (i.e. from the period [–30s, 0s] relative to the start of the regulation period at 0s) as follows:

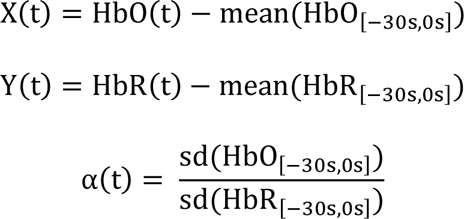

This preprocessed and motion-corrected signal *X*_corr_ (*t*) was then normalized relative to the HbO signal from the last five seconds of the no-regulation period (baseline) using the following formula:

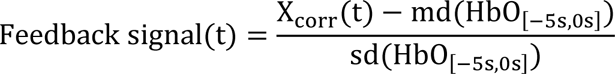

The feedback signal was further smoothed using linear interpolation over 1s. The final step was to convert the feedback value into visual feedback. This was implemented by mapping the feedback value onto a scale that ranged from 0 to 10, based on the following expression:

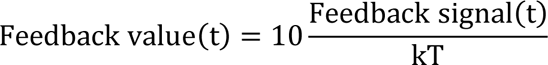

To receive positive feedback on downregulation, this value was multiplied by –1 for the downregulation group. In this expression, the maximum feedback value (level 10) was defined as a percentage (*k*) of a threshold value (*T*) that was determined for each participant based on their rTPJ activation during the reorienting of attention and perspective-taking tasks at pre-assessment. Similar to the calculation of the feedback signal during the neurofeedback task, the rTPJ activation during valid/invalid and perspective-taking/non-perspective-taking blocks was estimated and averaged over blocks. The mean between the contrasts of invalid vs valid (reorienting of attention task) and perspective taking vs non-perspective taking (vPT task) was calculated using the following formula:

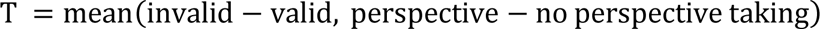

If this value was negative, only the positive contrast was used. If both contrasts were negative, the initial threshold was set to the default value *T =* 1. To scale the feedback signal, we defined level 10 of the feedback value as *k =* 0.25 (i.e., 25% of the threshold value *T*). The feedback value at each time was used to update the visual display in two ways. Each level of this scale from 0 to 10 was associated with eleven images of the avatar smiling to different extents with 10 being the largest smile. Therefore, the feedback value was used to update the image of the avatar displayed on the screen. Additionally, whenever the participant exceeded a feedback level of 5, a green frame appeared around the feedback display indicating a monetary reward (1 cent/sec).

To maintain the difficulty of the task across runs, the value of *k* was incremented by 0.25 for the next run if the participant exceeded level 5 for 75% of the time on each run.

### 2.5 Data processing and analysis

#### Data availability statement

Data of this study will be made available on a publicly accessible database after publication of the study as far as data protection regulations permit.

#### Statistical methods and software

The additional fNIRS offline analyses were carried out using Homer v2.8 (Huppert et al., 2009)(Huppert et al., 2009) and in-house Matlab scripts (Matlab 2018b; The Mathworks Inc, Natick, MA). Statistical analyses were performed using R (R Core Team, 2021). To assess the effects of the neurofeedback training, we calculated linear mixed models using the R packages lme4 (Bates et al., 2015) and lmerTest (Kuznetsova et al., 2017). The models were fitted using REML. In the case of non-normal residuals, robust nonparametric analysis of longitudinal data in factorial designs was carried out using the nparLD package (Noguchi et al., 2012), and ANOVA-type statistics (ATS) were reported. In addition, we calculated paired and Welch’s unequal variances t-test and Mann-Whitney/Wilcoxon tests for comparisons of mean values. Spearman’s rank correlation was used to assess relationships between neurofeedback regulation success, behavioral effects, and psychosocial factors. To explore predictors of behavioral improvements, we calculated stepwise multiple regression models and applied an Akaike information criterion (AIC) stepwise model selection algorithm (Akaike, 1974) to select the best models. Data are presented as means ± standard deviation (SD) unless indicated otherwise. For all analyses, a *p*-value below 0.05 was considered significant. Bonferroni correction was applied for the correlational analyses. We calculated Cohen’s d for mean comparisons or the correlation coefficient after non-parametric tests and partial eta-squared *(η_p_²)* for linear mixed models.

#### Neurofeedback regulation success

##### Further preprocessing

To analyze neurofeedback regulation success, we analyzed the time series of the feedback signal based on the online analysis adding further steps for artifact removal. First, we detected and automatically removed noisy channels by calculating coefficients of variation (CoV) and excluding channels with a CoV > 10% in HbO or HbR or channels with a variation difference between the chromophores of over 5%. In addition, channels in which we identified a flat line of at least 1s were removed (Bell et al., 2020). If the channel covering the rTPJ (COI) was detected as a noisy channel, we visually inspected the raw and preprocessed time series of the respective channel, and reincluded the channel if the high CoV was driven by spikes or drifts that could be removed by our preprocessing pipeline. The removed values were replaced by the average activation of the six neighboring trials. Second, outliers were removed if they exceeded 3SDs from the mean on-trial level and replaced by the last observation.

##### Additional robustness checks

Since short channel measurements were unfortunately not available for our system, we carried out an additional stepwise offline analysis approach to further test the robustness of the observed effects and to rule out the possibility that the neurofeedback signal change was driven by systemic physiological signals. For the offline robustness checks we used raw fNIRS signals of the same data sets as for the online analysis and carried out the same preprocessing and analysis steps as in the online analysis (bad channel removal, outlier detection, interpolation, bandpass filter, 5s moving average filter, and CBSI). In addition, we applied a more stringent bandpass filter (0.01-0.09 Hz), which is recommended by Pinti et al. (2019)Pinti et al. (2019) and should remove most of the systemic physiological signals (first robustness check). In the second, more conservative robustness check, we applied the common average reference (CAR) using the average time series of the 22 channels placed over the left hemisphere and subtracting it from the feedback channel time series. The CAR is considered to be a viable approach when short channel measurements are not available (Yücel et al., 2021), albeit a suboptimal one, since there is a risk of overcorrecting the signal or inducing additional effects depending on network activity during the task (Hudak et al., 2018; Kohl et al., 2020; Klein et al., 2022).

For all three analysis approaches, we calculated the median and standard deviation of the feedback signal time series for each of the 30s trials, normalized to the last five seconds of the no-regulation period (see formula for the calculation of the feedback signal). Mean values were calculated for each neurofeedback run and used for further group-level analyses.

##### Neurofeedback success measures

The main goal of the study was to test for successful control of rTPJ activation. However, there is no consensus on how to define neurofeedback regulation success (Paret et al., 2019; Kohl et al., 2020), and there is evidence of insufficient reporting quality in the field of fNIRS-based neurofeedback (Kohl et al., 2020), making it difficult to assess the effectiveness of a newly developed neurofeedback training protocol. Therefore, here we report several different neurofeedback success measures on the group and on the individual level, each of which has implications for the conclusion of the successful control of rTPJ activation (see Table 1). Besides measure of signal amplitude, we also include measures of signal variability in our analysis as they might be indicative of learning as well (see Kohl et al., 2020).

**Table 1.**
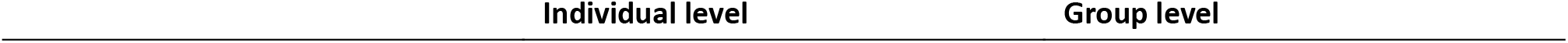
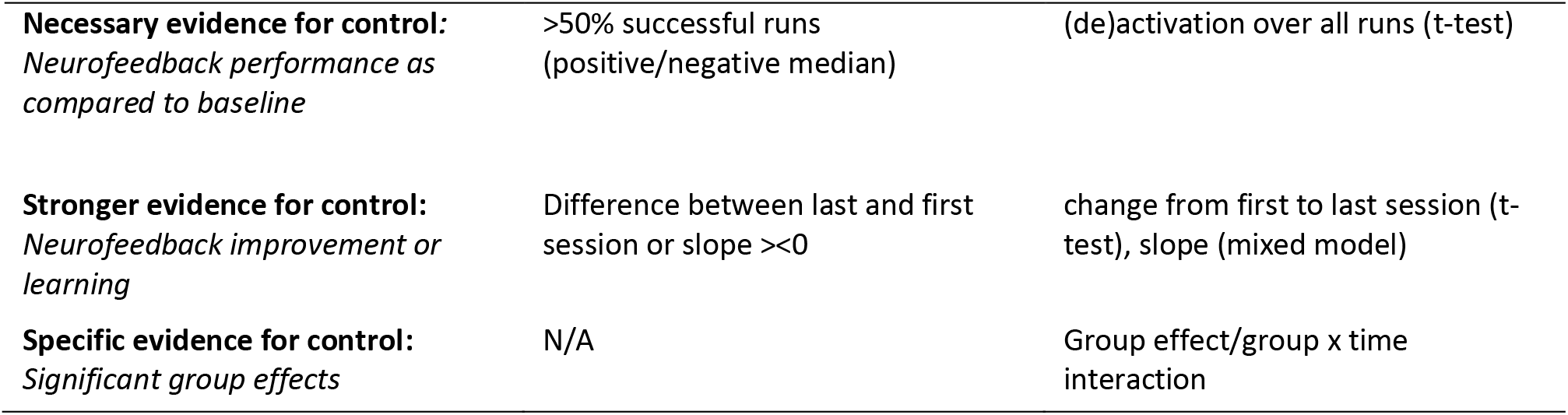
Neurofeedback success measures

First, we tested whether a participant is able to (1) activate the target region in the desired direction and maintain it over the course of the training compared to a within-baseline condition (*neurofeedback performance as compared to baseline – maintenance of activation*). On the group level, we used one-sample *t*-tests to test for a regulation effect against baseline for both groups separately. For the analysis on the individual level, we followed an exploratory approach according to Haugg et al. (2021) and Auer et al. (2015) to classify successful participants who maintained up- or downregulation throughout the training, independent of a learning effect. A run was classified as successful if the medians of the feedback signal over trials were positive in the upregulation group and negative in the downregulation group. Participants who demonstrated more than 50% successful neurofeedback runs were then classified as “successful”, and participants below 50% as “unsuccessful”. The numbers of successful participants are reported. In addition, we report on the numbers of successful runs per participant (see Table S1 and S2).

These measures only provide *necessary evidence for the control* of baseline brain activation, but they alone are insufficient to draw conclusions about the successful regulation of brain activation through neurofeedback. This limitation arises because the activation of the target region could be influenced by factors inherent to the experimental paradigm, such as stimulation from the experimental stimuli, or the use of mental strategies, rather than solely attributable to the effects of neurofeedback.

*Stronger evidence for control* would be if a participant showed a voluntary change in (2) *amplitudes* and (3) *variability* of the feedback signal over time compared to a within-baseline condition (*neurofeedback improvement* or learning). On the group level, we tested for a time effect of rTPJ regulation using linear mixed models or non-parametric ANOVAs for both groups separately. On the individual level, the neurofeedback improvement of each participant was calculated based on the slope of the linear regression over all neurofeedback runs. Here, a participant was classified as “successful” if he or she showed a slope larger than 0 in the upregulation group and smaller than 0 in the downregulation group. In addition, since learning does not necessarily follow a linear trajectory, we compared the regulation success of the last session with the first session. On the group level, we used the paired *t*-test for both groups separately to compare rTPJ activation in the last session compared to the first session. On the individual level, we classified a participant with a positive/negative value as “successful” and vice versa.

Lastly, we tested for a specific effect of regulation *(specific evidence for control)* by comparing measures 1-3 with the between-group control condition. To do so, we calculated linear mixed models or non-parametric ANOVAs and tested for a significant group effect (1) and a significant group × time interaction (2-3).

#### Behavioral effects

For the reorienting of attention task, only correct RTs were analyzed. RTs <100 ms and >1000 ms, as well as incorrect key presses were excluded from the analysis. Harmonic means of valid and invalid trials of the invalid blocks were calculated and analyzed. The harmonic mean, as recommended for RT analysis by Ratcliff (1993), is a more unbiased estimator of the central tendency of RTs than the arithmetic mean, which also reduces the effects of outliers while remaining high power. In addition, RTs for invalid trials were subtracted from valid trials to estimate the costs of shifting attention from the cued position to a non-cued target (reorienting effect). Two participants, one from each group, had to be excluded due to a technical error or not understanding task instructions. For the vPT task, harmonic means of the RTs and mean accuracies were analyzed. One participant had to be excluded from this analysis due to a technical error. For both tasks, linear mixed models or non-parametric ANOVAs were calculated with the task condition and measurement time as within-subject factors and the group as the between-subject factor. According to our hypotheses, we expected to see a significant group × time interaction as well as a significant within-group time effect in the upregulation group for both tasks.

To further confirm the specificity of the behavioral effects and control for unspecific contributions of psychosocial factors, we calculated four sets of correlational analyses:

1-2) Observed behavioral effects in the attention task/vPT task were correlated with three different neurofeedback success measures for both groups separately as well as across group resulting in 27 tests (3 conditions x 3 success measures x 3 groups) for each task.

3-4) Changes in RTs of the attention task/changes in accuracies of the vPT task across conditions were correlated with eleven results of questionnaires assessing psychosocial factors (e.g., expectations toward the training, subjective evaluation of the training, etc.) resulting in 33 tests (11 questionnaire results x 3 groups) for each task.

We applied the Bonferroni correction separately for the four different sets of analyses, each involving a distinct number of tests (i.e., 27, 27, 33, and 33).

#### Predicting behavioral improvements

As neurofeedback represents a potentially useful tool for application in clinical populations exploring how subclinical symptoms, personality traits, and baseline task performance are related to specific behavioral neurofeedback, effects in healthy samples can inform clinical translation. In terms of TPJ functioning, these include ASD symptoms (Kana et al., 2014, 2015, 2016) and measures of empathy as well as baseline cognitive and socio-cognitive performance data.

We only found a specific effect in the reorienting of attention task and no specific effect in the vPT task. Therefore, we conducted an analysis for the effects in the reorienting of attention task using absolute ΔRTs across conditions as a dependent variable of a multiple regression model. We used the results of questionnaires assessing autism-related traits and empathy (AQ, EQ, SQ, SRS, IRI) as well as baseline task performance (RTs across conditions of the reorienting of attention task and accuracies in PT trials of the vPT task) as predictor variables (7 in total). To avoid overfitting, stepwise multiple regression models were calculated and the AIC stepwise model selection algorithm (Akaike, 1974) was used to select the best model.

#### Mental strategies underlying neurofeedback regulation

Based on a content analysis of the strategy reports, we identified 20 different categories of strategies that participants employed to regulate their brain activity. Figure 8 in the results section shows the different categories and their distribution. We classified the reported strategies into the different categories, calculated how many strategies were used by each subject and how many participants reported to have used a particular strategy. The mean number of strategies used, and the frequencies of the different strategies were compared between groups.

## 3 Results

### 3.1 Baseline characteristics

There were no baseline differences between the two groups, i.e., neither in the questionnaire data nor in the reaction times and accuracies in the (1) reorienting of attention task and (2) vPT task (all *p* > 0.05; see Table 2, Figure 6, and Table S1-2 for more detailed baseline characteristics and questionnaire results). In addition, the thresholds for the feedback signal as determined by rTPJ activation during the pre-assessments did not significantly differ between the groups (upregulation group = 2.19 ± 1.45, range 0.45 – 6.2, downregulation group = 2.76 ± 1.84, Range 0.03 – 6.92). These results demonstrate that our randomization procedure was successful. Seven participants (5 in the upregulation and 2 in the downregulation group) showed negative contrasts for both tasks.Therefore, their initial threshold was set to a default value of 1.

**Table 2.** Baseline characteristics and questionnaire results

|  | Upregulation<br>(*M ± SD*) | Downregulation<br>(*M ± SD*) | *p-value* |
| --- | --- | --- | --- |
| N | 27 (13 female) | 18 (9 female) |  |
| Age (years) | $24.22 \pm 3.03$ | $24.22 \pm 2.71$ | 0.935 |
| pre RTs attention task – invalid | $497 \pm 69$ ms | $508 \pm 90$ ms | 0.816 |
| pre RTs attention task – valid | $452 \pm 61$ ms | $468 \pm 94$ ms | 0.703 |
| pre accuracies attention task - invalid | $0.98 \pm 0.02$ | $0.98 \pm 0.05$ | 0.169 |
| pre accuracies attention task - valid | $0.99 \pm 0.02$ | $0.99 \pm 0.03$ | 0.112 |
| pre RTs vPT task – PT | $3667 \pm 327$ ms | $3630 \pm 302$ ms | 0.701 |
| pre RTs vPT task – NPT | $3611 \pm 286$ ms | $3620 \pm 272$ ms | 0.966 |
| pre accuracies vPT task – PT | $0.946 \pm 0.071$ | $0.926 \pm 0.091$ | 0.634 |
| pre accuracies vPT task – NPT | $0.981 \pm 0.035$ | $0.971 \pm 0.044$ | 0.230 |
| pre rTPJ thresholds | $2.19 \pm 1.45$ | $2.76 \pm 1.84$ | 0.270 |
| AQ total | $15.30 \pm 6.14$ | $13.94 \pm 4.49$ | 0.399 |
| EQ total | $45.19 \pm 9.76$ | $45.17 \pm 8.28$ | 0.995 |
| SQ total | $29.96 \pm 10.35$ | $32.11 \pm 13.75$ | 0.577 |
| IRI total<sup>1</sup> | $56.78 \pm 11.78$ | $52.39 \pm 8.83$ | 0.161 |
| SRS: total | $39.26 \pm 19.36$ | $37.94 \pm 12.94$ | 0.862 |
| POMS: depression/anxiety | $0.40 \pm 0.52$ | $0.42 \pm 0.58$ | 0.716 |
| POMS: vigor | $3.43 \pm 1.01$ | $3.40 \pm 0.90$ | 0.920 |
| POMS: fatigue | $1.61 \pm 0.99$ | $1.53 \pm 1.12$ | 0.814 |
| POMS: hostility | $0.54 \pm 0.93$ | $0.52 \pm 0.77$ | 0.737 |
| General self-efficacy | $30.59 \pm 2.50$ | $31.83 \pm 3.57$ | 0.211 |
| Expectations | $2.42 \pm 0.56$ | $2.67 \pm 0.59$ | 0.123 |
| Motivation | $3.70 \pm 0.40$ | $3.63 \pm 0.50$ | 0.695 |

<sup>1</sup> according to Cliffordson (2001) and Paulus (2012) Cliffordson (2001) and Paulus (2012); AQ, Autism Spectrum Quotient; ASRS, Adult ADHD Self-Report Scale; EQ, Empathy Quotient; IRI, Interpersonal Reactivity Index; POMS, Profile of Mood States; PT, perspective taking; NPT, non-perspective taking; SQ, Systemizing Quotient; SRS, Social Responsiveness Scale; vPT, visual perspective taking.

### 3.2 Regulation behavior and rewards

Both groups were able to regulate the signal in the desired direction and remained above their individual threshold. On average, the upregulation group remained above the threshold in each trial for a longer period (*M* = 16±3.25 of 30s) than the downregulation group (*M* = 9.41±2.9 of 30s), but only the downregulation group improved over time (see Supplementary Material 3.1 and Figure S1 for detailed results). As a result, the upregulation group also received significantly more monetary rewards than the downregulation group (*M* = 12.80±2.33€ vs *M* = 7.86€±2.16€; *t*(38,4) = 7.29, *p* < 0.001, *d* = 2.35).

### 3.3 Neurofeedback regulation success

Table 3 shows the results for the different neurofeedback success measures and Figure 5 shows grand averages of HbO changes of the feedback signal for all four neurofeedback training days (sessions) and box plots of average feedback performance based on the online analysis for all twelve neurofeedback runs. Tables S5-6 show the individual results of neurofeedback regulation success.

**Figure 5.**
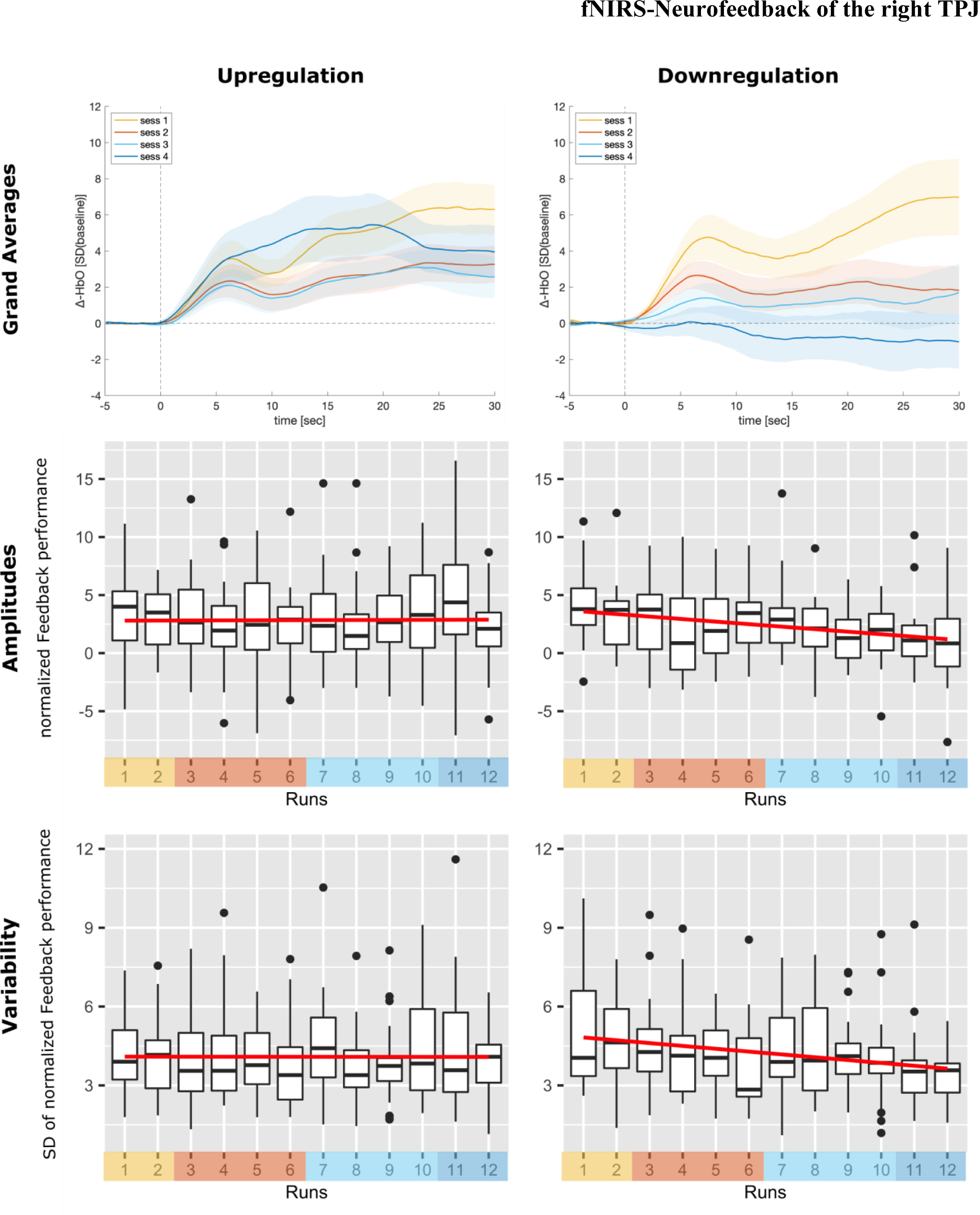
Neurofeedback regulation performance. The first row shows the grand averages of the changes in HbO of the feedback channel for the four neurofeedback training days (sessions). The second row shows box plots of the average feedback performance as assessed by the standardized median change of rTPJ activation averaged over participants for all twelve neurofeedback runs (sessions color-coded) based on the online analysis. The third row shows box plots of the standard deviations of feedback performance as assessed by the standardized median change of right TPJ activation averaged over participants for each run. The regression lines of the linear models are depicted in red. Paired-sample t-tests comparing the last session (run 11 and 12) with the first session (run 1 and 2) revealed significant effects and the ANOVA over all neurofeedback runs only revealed non-significant time trends in the downregulation group only. No time effect was observed in the upregulation group (see main text).

**Table 3.** Neurofeedback regulation success

|  | Online analysis (amplitudes) |  |  | Online analysis (variability) |  |  |
| --- | --- | --- | --- | --- | --- | --- |
| Upregulation group | *M*±*SD* | *n* | *p*-value | *M*±*SD* | *n* | *p*-value |
| NF performance - compared to baseline | 2.84±2.2 | 26/27 | < 0.001* | N/A | N/A | N/A |
| NF improvement (slope) | 0.01±0.34 | 14/27 | 0.84 | 0.00±0.17 | 17/27 | 0.595 |
| NF improvement (last vs first) | 0.02±3.7 | 15/27 | 0.97 | -0.06±1.69 | 17/27 | 0.86 |
| Downregulation group | *M*±*SD* | *n* | *p*-value | *M*±*SD* | *n* | *p*-value |
| NF performance - compared to baseline | 2.39±1.89 | 4/18 | 1 | N/A | N/A | N/A |
| NF improvement (slope) | -0.22±0.35 | 15/18 | 0.09 | -0.11±0.17 | 15/18 | 0.143 |
| NF improvement (last vs first) | -2.36±3.6 | 15/18 | 0.01* | -1.37±1.76 | 15/18 | 0.004* |

Neurofeedback regulation success according to different success measures for both groups. The p-values reflect the results of the group analysis as described in 2.5 “Data processing and analysis”; NF, neurofeedback.

#### Regulation success (amplitudes)

In the upregulation group, we observed high rTPJ activation that was sustained over the course of the training. In contrast, the downregulation group unexpectedly showed the same effect (activation instead of deactivation), which, however, disappeared over the course of the training. One-sample *t*-tests revealed a significant main effect of regulation over all runs in the upregulation group (*M* = 2.84±2.2, *t*(26) = 6.72, *p* < 0.001, *d* = 1.32) and the downregulation group (*M* = 2.39±1.89, *t*(17) = 5.6, *p* < 0.001, *d* = 1.36), meaning that on average, rTPJ activity also increased in the downregulation group. Paired-sample *t*-tests, however, only revealed a significant decrease between the last and the first session in the downregulation group (*Mdiff* = −2.36±3.6, *t*(17) = 2.79, *p* = 0.01, *d* = 0.68), but no significant increase in the upregulation group (*Mdiff* = 0.02±3.7, *p* > 0.98, *d* = −0.01). The non-parametric ANOVA only revealed a non-significant time trend in the downregulation group (*F_ATS_* (5.85, ∞) = 1.86, *p* = 0.09) and no effect in the upregulation group. No specific group effect or significant group × time interaction was found.

The analysis on the individual level revealed that in the upregulation group, 96.30% of the participants (26 of 27) were successfully upregulating rTPJ activity (>50% successful runs; *M* = 9.63 ± 2.27), 51.85% (14 of 27) showed an improvement of regulation performance over runs as indicated by a positive slope, and 55.56% (15 of 27) showed a higher regulation performance in the last session compared to the first session. In the downregulation group, only 22.22 % (4 of 18) were successfully downregulating rTPJ activity (>50% successful runs; *M* = 3 ± 2.66), 83.33% (15 of 18) showed an improvement of regulation performance over runs as indicated by a negative slope and a higher regulation performance in the last compared to the first session.

#### Regulation success (variability)

For the variability of the neurofeedback performance over time, similar results compared to the main analysis of regulation success (analysis based on signal amplitudes) were observed. Paired-sample *t*-tests also revealed a difference between the last and the first session in the downregulation group (*Mdiff* = −1.37±1.76, *t*(17) = 3.29, *p* = 0.004, *d* = 0.8) but not in the upregulation group (*Mdiff* = −0.06±1.69, *p* = 0.86, *d* = 0.03). The non-parametric ANOVA only revealed a non-significant time trend in the downregulation group (*F_ATS_* (7.27, ∞) = 1.55, *p* = 0.143). No specific group effect or significant group × time interaction was found.

The individual analysis revealed that in the upregulation group, 62.96% of the participants (17 of 27) showed decreasing standard deviations over runs, as indicated by a negative slope of the regression and lower values in the last session compared to the first session. On the other hand, in the downregulation group, 15 out of 18 participants (83.33%) showed decreasing standard deviations over runs, as indicated by a negative slope of the regression and lower values in the last session compared to the first session.

#### Robustness checks

Robustness check 1 successfully confirmed the results of the online analysis. However, none of the effects survived the more conservative robustness check 2 (see Supplementary Material 3.3 for detailed results).

### 3.4 Primary behavioral outcomes

#### Reorienting of attention task

As expected, we found a significant main effect of condition for RTs (*F*(1,123) = 111.21, *p* < 0.001, *η_p_²* = 0.47), and accuracy data (*F_ATS_* (1, ∞) = 17.18, *p* < 0.001), reflecting a significant reorienting effect in both groups across time points (*mean RTs valid* = 456 ± 77ms, *mean RTs invalid* = 499 ± 80ms). The hypothesized three-way interaction of group × time × condition was not observed, i.e., no significant time effects or group × time interaction effects were observed for the reorienting effect. However, we found a significant group × time interaction (*F*(1,123) = 17.17, *p* < 0.001, *η_p_²* = 0.12), and a significant main effect of time in both groups (upregulation group: (*F_ATS_* (1, ∞) = 6.20, *p* = 0.013), downregulation group: (*F_ATS_* (1, ∞) = 4.42, *p* = 0.036), indicating a group-specific effect of the training on RTs across conditions. The pre-post comparisons revealed that after the neurofeedback training, reaction times across conditions decreased in the upregulation group (*pre* = 474 ± 68ms, *post* = 457 ± 57ms, *d* = 0.51) and increased in the downregulation group across conditions (*pre* = 488 ± 93ms, *post* = 503 ± 108ms, *d* = −0.56). No other main effects or interactions were found (see Figure 6).

**Figure 6.**
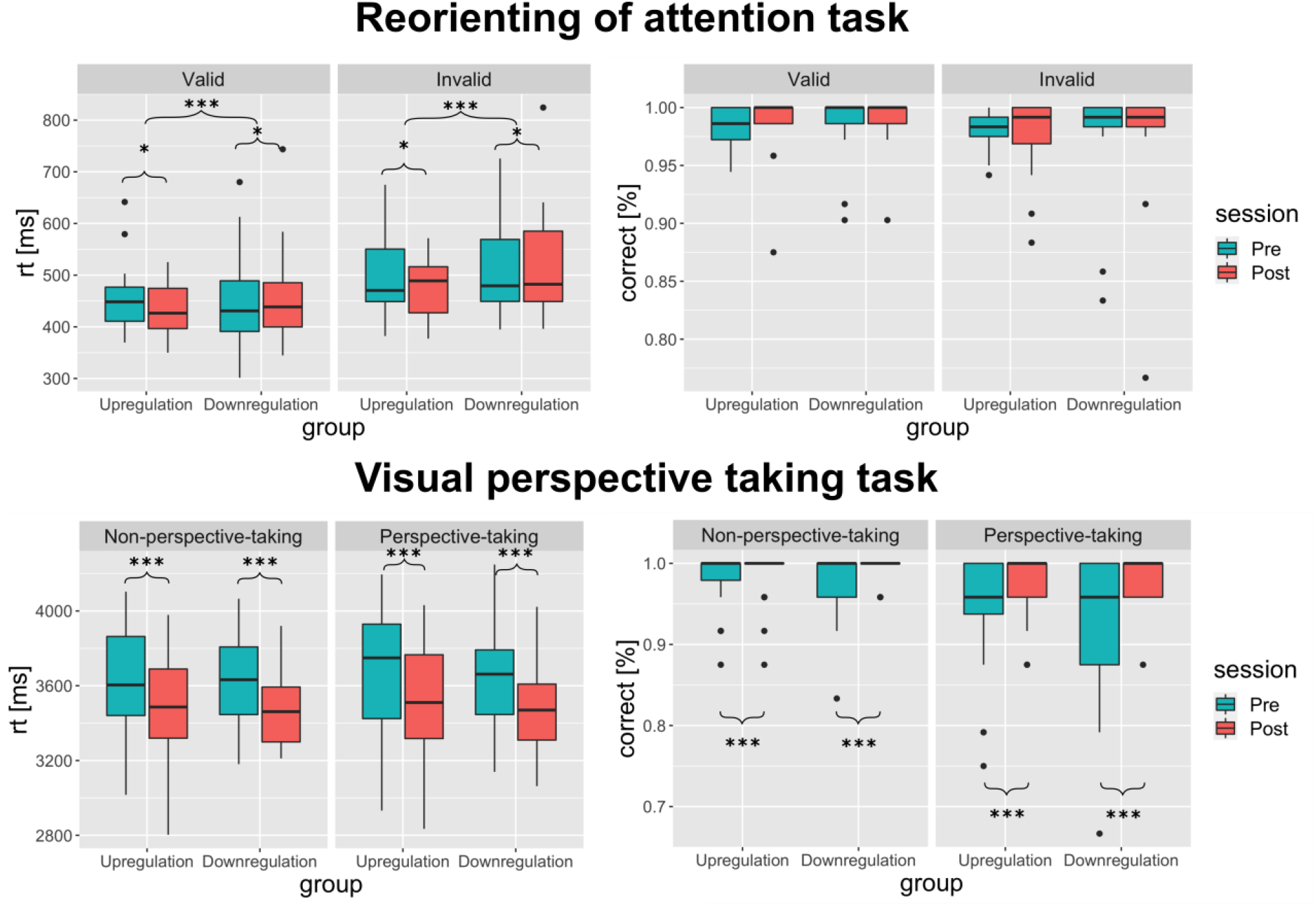
Primary behavioral outcomes. Results of the reorienting of attention task (upper panel) and visual perspective-taking task (lower panel). For detailed descriptive statistics, see Table S2. Boxplots show interquartile range ± 1.5 (whiskers). Asterisks denote the significance for the group × time interaction and within-group time effects across task conditions; *** p < 0.001; * p < 0.05.

If we included trials from the valid only blocks, results did not change, but the time effect in the downregulation group (*F_ATS_* (1, ∞) = 3.25, *p* = 0.071) failed to reach significance (see Supplementary Material 4).

#### vPT task

Contrary to our hypothesis, the three-way interaction of group × time × condition was neither observed for RTs nor for accuracies in the vPT task. RTs decreased in both groups (*F*(1,126) = 55.58, *p* < 0.001, *η_p_²* = 0.31) irrespective of condition (pre = 3630 ± 270ms, post = 3500 ± 295ms, *d* = 0.83). No other main effects or interactions were significant. Accuracies increased in both groups (pre = 95.8 ± 3.1%, post = 98.3 ± 6.5%, *d* = 0.58), as indicated by a significant time effect (*F_ATS_* (1, ∞) = 11.91, *p* < 0.001) and a significant condition effect (*F_ATS_* (1, ∞) = 10.75, *p* < 0.005), but no interaction effect occurred.

However, a ceiling effect was observed in this task. The majority of the participants responded with 100% accuracy in this task during the pre-assessment (29 in the NPT and 18 in the PT condition) and during the post-assessment (34 in the NPT and 29 in the PT condition; see Figure 6)).

### 3.5 Mental strategies, secondary outcomes, and unspecific psychological effects

#### Mental strategies underlying neurofeedback regulation

The downregulation group used significantly more different strategies (*M* = 8.66±2.47) during the neurofeedback training compared to the upregulation group (*M* = 6.26±3.24; *t*(42.11) = 2.82, *p* = 0.007, *d* = 0.87). Figure 7 shows the distribution of strategies as reported by the participants of both groups. Fisher’s exact Chi-square test revealed no significant association between the group and reported strategies (*p* = 0.982), indicating that similar strategies were used for both upregulating and downregulating TPJ activity. Table S8 shows the percentages of strategies relative to the total number of strategies reported per group and their mean success rating. In total, most strategies were reported to be more successful in the upregulation group (mean success rating: 3.35) than in the downregulation group (mean success rating: 2.74), and socio-cognitive strategies and positive mental imagery were reported most frequently in both groups (see Supplementary Material 5).

**Figure 7.**
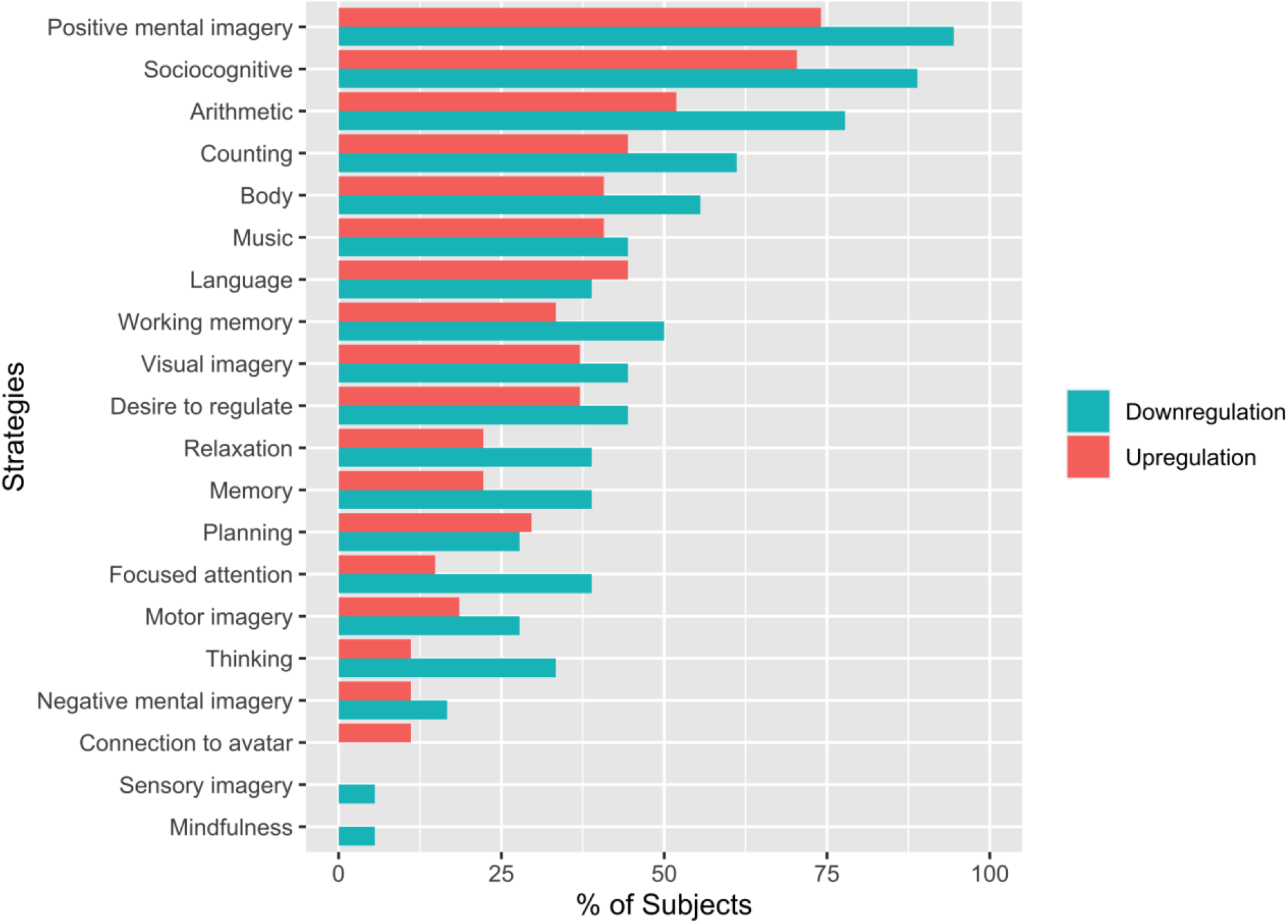
Strategies as reported by participants for each group.

#### Motivation, self-control beliefs, self-efficacy, and mood

Motivation to take further part in the neurofeedback training as assessed after each session was high in both groups (*Mdn* = 9, on a 10-point rating scale), but decreased slightly in the upregulation group over the course of the sessions. There was a significant time effect (*F_ATS_* (2.34, ∞) = 3.11, *p* = 0.04), which was driven by a simple main effect of time in the upregulation group (*F_ATS_* (2.54, ∞) = 3.48, *p* = 0.02). No time effect was observed in the downregulation group. Post hoc comparisons of the last session with the first session confirmed a slight, but significant, decrease of motivation in the upregulation group (first session, *Mdn* = 9, last session, *Mdn* = 8, *p* = 0.02, *r* = 0.491) and no effect in the downregulation group (first session, *Mdn* = 9.5, last session, Mdn = 9.5, *p* = 0.43, *r* = 0.124; see Table S4).

For the general self-efficacy scale, we found a significant time effect (*F*(1,43) = 4.93, *p* = 0.03, *η_p_²* = 0.10). Although the group × time interaction failed to reach significance (*F*(1,43) = 2.39, *p* = 0.129, *η_p_²* = 0.05), this effect seemed to be driven by an increase in the upregulation group from pre- (*M* = 30.59 ± 2.5) to post-assessments (*M* = 32.04 ± 3.16). This was indicated by a simple main effect of time, which, however, failed to reach significance (*F*(1,52) = 3.48, *p* = 0.07, *η_p_²* = 0.06). No time effect was observed in the downregulation group (*F*(1,34) = 0.02, *p* = 0.89, *η_p_²* = 0; see Table S3).

Participants’ beliefs of how well they could control the neurofeedback signal was lower in the downregulation group at the beginning of the training, but increased to the level of the upregulation group towards the end of the training, as indicated by a significant time effect (*F*(3,128.3) = 3.36, *p* = 0.02, *η_p_²* = 0.07), group effect (*F*(1,42.98) = 11.88, *p* = 0.001, *η_p_²* = 0.26) as well as a significant group × time interaction (*F*(3,128.2) = 6.17, *p* < 0.001, *η_p_²* = 0.12). A simple main effect of time was only observed in the downregulation group (*F*(3,68) = 4.75, *p* = 0.005, *η_p_²* = 0.15). Post hoc t-tests indicated that there was a group difference in the first neurofeedback session (upregulation group = 7.19±1.42, downregulation group = 4.44±1.82, *t*(30.53) = 5.37, *p* < 0.001, *d* = 1.95) and the second neurofeedback session (upregulation group = 6.92±1.5, downregulation group = 5.72±1.64, *t*(34.46) = 2.48, *p* = 0.02, *d* = 0.87), which disappeared in the third session (upregulation group = 7.08±1.35, downregulation group = 6.56±1.82) and the last session (upregulation group = 6.88±1.82, downregulation group = 6.36±2.11) see Table S4). The neurofeedback training showed no significant effect on mood states, as assessed with the POMS (see Table S3).

#### Expectations and evaluations of the training

No differences were found between the groups with respect to the expectation towards the neurofeedback training and the subjective evaluation of the training (believed efficacy, joy, and experimenter). The debriefing questionnaires revealed, however, that 71.11% of the participants (80.77% in the upregulation and 55.56% in the downregulation group) guessed the group assignment correctly, although most participants reported that they were not confident about their judgement.

### 3.6 Correlations of behavioral outcomes with regulation performance and psychosocial factors

We found a significant negative correlation between changes in RTs in the valid trials of the reorienting of attention task and neurofeedback performance, as assessed by the number of successful runs (*rho* = −0.47, *p* = 0.045, Bonferroni corrected), indicating higher improvements of RTs in participants with more successful runs in both groups. Subgroup analysis revealed no significant effect after Bonferroni correction.

For the perspective-taking task, we found a significant correlation between neurofeedback improvement (slopes) and improvements in the accuracies of NPT trials across groups (*rho* = −0.49, *p* = 0.02), indicating greater performance improvements in participants who were more successful in learning downregulation over the course of the training. This significant correlation was only observed in the downregulation group (*rho* = −0.71, *p* = 0.039, Bonferroni corrected).

None of the psychosocial factors correlated significantly with behavioral outcomes after Bonferroni correction. For more details including significant correlations on the uncorrected level, see Supplementary Material 6.

### 3.7 Predicting behavioral improvements

For the neurophysiologically specific improvements observed in the attention task, we found that IRI total scores and baseline performance in the attention task predicted changes in performance (see Table 4). The subgroup models revealed that baseline attentional performance and EQ scores only predicted behavioral improvements in the upregulation group, thus indicating greater improvements in participants with lower baseline performance and higher EQ scores. In the downregulation group, IRI scores and baseline vPT performance predicted decreased performance in the attention task after the training.

**Table 4.** Summary statistics of the stepwise multiple linear regression model predicting behavioral improvements.

| **Model summary – both groups** |  |  |  |  |
| --- | --- | --- | --- | --- |
| *R<sup>2</sup>* | *Adjusted R<sup>2</sup>* | *Residual SE* | *F(2,39)* | *p-value* |
| 0.377 | 0.346 | 0.022 | 11.82 | <0.001*** |
| **Coefficients** |  |  |  |  |
| *Step* | *Beta* | *SE* | *t-value* | *p-value* |
| Intercept | -0.102 | 0.027 | -3.836 | <0.001*** |
| IRI total | 0.001 | 0.0004 | 2.143 | 0.038* |
| pre RTs attention task | 0.17 | 0.047 | 3.651 | <0.001*** |
| **Model summary – Upregulation group** |  |  |  |  |
| *R<sup>2</sup>* | *Adjusted R<sup>2</sup>* | *Residual SE* | *F(2,23)* | *p-value* |
| 0.513 | 0.471 | 0.026 | 12.12 | <0.001*** |
| **Coefficients** |  |  |  |  |
| *Step* | *Beta* | *SE* | *t-value* | *p-value* |
| Intercept | 0.169 | 0.039 | 4.308 | <0.001*** |
| EQ | -0.001 | 0.001 | -2.311 | 0.03* |
| pre RTs attention task | -0.271 | 0.086 | -3.170 | 0.004** |
| **Model summary – Downregulation group** |  |  |  |  |

| *R*<sup>2</sup> | *Adjusted R*<sup>2</sup> | *Residual SE* | *F*(2,13) | *p*-value |
| --- | --- | --- | --- | --- |
| **0.463** | 0.38 | 0.023 | 5.596 | 0.018* |
| **Coefficients** |  |  |  |  |
| *Step* | *Beta* | *SE* | *t*-value | *p*-value |
| **Intercept** | -0.403 | 0.13 | -3.106 | 0.008** |
| **IRI total** | 0.003 | 0.001 | 2.752 | 0.017* |
| **pre accuracy vPT task** | 0.286 | 0.108 | 2.634 | 0.021* |

Note that we used absolute DRTs in the reorienting of attention task in the model including both groups but the real DRTs in the subgroup models.

## 4 Discussion

This is the first study demonstrating the feasibility and effectiveness of neurofeedback training of the rTPJ based on fNIRS. We demonstrated successful activation of the rTPJ *compared to baseline* (*necessary evidence for control*) within the first training session (2 neurofeedback runs) in the upregulation group. Only one of 27 participants in this group failed to activate (<50% successful trials). However, we observed no significant effect of *neurofeedback improvement*; almost half of the participants (13 of 27) failed to show a positive slope. Successful downregulation, on the other hand, required at least four sessions (12 neurofeedback runs) or more. Most participants failed to successfully downregulate, but a significant *neurofeedback improvement* effect was observed in this group and only three of 18 participants failed to show such an effect. Surprisingly, participants in the downregulation group were also activating their rTPJ at the beginning of the training but learnt to downregulate or at least to not activate it anymore towards the end of the training. This can be interpreted as *strong evidence for control* in the downregulation group.

While only unspecific improvements were observed for vPT, specific up/down-regulatory effects on stimulus-driven attention were observed in the reorienting of attention task, providing evidence for a neurophysiological specific effect of rTPJ regulation on stimulus-driven spatial attention, although not specifically related to the reorienting process of attention (as indicated by a reduced invalidity effect). Neurophysiological specificity was further confirmed by the fact that non-specific psychological mechanisms and mental strategies did not differ between groups and therefore cannot explain the group effect. The training was well received by the young and healthy participants with no dropouts as well as high levels of motivation and feelings of control reported throughout the training.

### 4.1 Neurofeedback regulation success

While we demonstrate the feasibility and effectiveness of a neurofeedback training of the rTPJ, the specific results of different neurofeedback success measures, in conjunction with the findings of the behavioral effects, yield a complex picture.

As both groups showed high activation of the rTPJ from the beginning of the training and only the downregulation group showed a learning effect we cannot derive definitive conclusion regarding the effectiveness of a neurofeedback upregulation training.

The initial high activation of the rTPJ might be explained by the contribution of general neurofeedback regulation mechanism by a neurofeedback controller network. Such a separate controller network involves neural populations of the TPJ (Emmert et al., 2016; Sitaram et al., 2017) possibly related to the integration of visual feedback as well as other feedback-related processes such as prediction processing (Bzdok et al., 2013). Therefore, there might have been an overlap of the neural populations of the neurofeedback controller network with the neurofeedback target region, meaning that the measured activity at rTPJ could have been a combination of the two. This potential overlap complicates the interpretation of activity changes and the relevance of feedback. One may speculate that during the regulation period, the controller network initially increased activity at rTPJ, but extended learning led to changes in the network, potentially reducing its activity and leading to complex effects on the measured upregulation and downregulation conditions. In such a scenario decreased activity in the controller network counteracted a potential increase over time in the target region, diminishing an observable learning effect. While the baseline period served as a control for stimuli-evoked activity, it did not account for baseline activity specifically related to the controller network, which was only engaged during the regulation period. However, we acknowledge the speculative nature of this account, which can only be confirmed through fMRI studies employing more fine-grained measures and estimations of the neurofeedback controller network.

Furthermore, the complexity of the social neurofeedback stimuli as well as the instructions used in our design may explain the initial high activation of the rTPJ. The rTPJ involves parts of the posterior STS, an area which has been attributed to the face processing network, and a subregion of the STS closely located to anterior parts of the rTPJ which has been associated with biological motion as well as emotional face processing (Beauchamp, 2015; Müller et al., 2018). Although we used digitizer measurements to ensure the correct placements of the feedback channels over anterior parts of the rTPJ, given the spatial resolution of fNIRS and the variability in optode placements we cannot exclude the possibility that the feedback channel captured the activation of this subregion of the STS – at least in some of the participants. The activation of the feedback channel might therefore have been partly induced by the feedback stimuli when the avatar started smiling or even by participants paying more attention to the facial stimuli during the regulation condition.

Lastly, this effect might be explained by the fact that both groups received the same strategy instructions and as a result relied heavily on socio-cognitive strategies associated with rTPJ activation (Bzdok et al., 2013).

The absence of a learning effect in the upregulation group made it also difficult to detect, and may explain, the absence of a significant group x time interaction effect in the current study (*specific evidence for control)*.

Nevertheless, the observed neurofeedback learning effect in the downregulation group, along with the specific effects on the behavioral level, provides interesting and encouraging findings as they indicate a neurophysiologically specific mechanism of rrTPJ regulation on stimulus-driven attention. These results have important implications for future study designs and clinical translation we discuss below (see section 4.2 and 4.5).

The first robustness check further confirmed the results of the online analysis, but the second, more conservative robustness check did not. This should be interpreted with caution, since given the limited spatial resolution and coverage in our study the CAR approach involves the risk of overcorrecting the signal or inducing additional effects depending on network activity during the task (Hudak et al., 2018; Kohl et al., 2020; Klein et al., 2022). In particular, the first approach (bandpass filter of 0.01-0.09 Hz) is capable of removing most of the frequencies associated with systemic physiology, including heart rate (∼1Hz), breathing rate (∼0.3Hz), and Mayer waves (∼0.1Hz; Pinti et al., 2019). Moreover, we took care to keep the contribution of systemic physiological changes in our experimental paradigm at a low level by using variable stimulus onsets and instructing our young and healthy participants to calm down before the experiment, breathe regularly, and avoid unnecessary movements. Therefore, we can assume that it is very unlikely that the observed effects were driven by systemic physiology, but instead by the real neural activation of the rTPJ.

### 4.2 Primary behavioral outcomes

The upregulation group showed increased performance and the downregulation group decreased performance in the reorienting of attention task across conditions. Our single-blinded, bidirectional-regulation control group design allowed us to properly control for neurofeedback non-specific or general non-specific effects and demonstrate neurophysiological specificity of the observed behavioral effects. It is unlikely that differences in the employed strategies can explain the behavioral effects, since although the downregulation group showed a higher variation of strategies, both groups relied on the same strategies to regulate their brain activity. The absence of group-specific correlations of regulation success with behavioral outcomes and the missing group-specific effect in the vPT task, together with other non-significant correlations between behavioral improvements and psychosocial factors, underline the role of other non-specific psychosocial mechanisms such as reward, control beliefs, and expectations in explaining the behavioral effects. However, given the strength of our study design, the observed dissociation in the reorienting of attention task provides evidence for a specific neurophysiological effect of rTPJ regulation on stimulus-driven attention.

Contrary to our hypothesis, we did not find a specific effect on the reorienting of attention, i.e., a specific improvement in invalid trials in the upregulation group. This is surprising given the assumed specific role of the rTPJ in reorienting of attention (Krall et al., 2015), which has been supported by previous neurostimulation studies (Roy et al., 2015; Krall et al., 2016). However, neurofeedback is different from neurostimulation. Instead of passively receiving neurostimulation, neurofeedback training requires the active participation of the participant and the skill of neural regulation to be successfully learned, which involves the recruitment of additional neural networks throughout the training (Emmert et al., 2016; Sitaram et al., 2017) that may have induced additional behavioral effects. Possible downstream effects of rTPJ regulation on other brain activities may have also resulted in additional behavioral effects (Kvamme et al., 2022). Furthermore, the specific role of the rTPJ in early stimulus-driven attentional reorienting has been questioned by Geng and Vossel (2013), who suggest a rather general role in post-perceptual contextual updating and adjustments of top-down expectations.

We found increased RTs after downregulation training indicating decreased performance in stimulus-driven attention, which is an interesting albeit preliminary finding. To date, only a few studies have applied a bidirectional control group approach and demonstrated group-specific changes, also including decreases in performance, for example sustained attention and response inhibition in the study of Yamashita et al. (2017). Future studies should test the robustness of these effects, assess potential long-term effects, and explore if decreased performance after downregulation can also be observed in other cognitive domains. If so, the bidirectional control group approach would be an interesting tool for cognitive neuroscience studies, since it is more efficient and the demonstration of such a dissociation provides stronger (causal) evidence than just an upregulation effect. However, caution is advised with respect to long-term effects and when such designs are applied to clinical populations.

Only unspecific improvements were found for vPT. The observed improvement might have been the result of a retest effect or a ceiling effect, which was observed for most of the participants and may have masked a group-specific effect in this task. Beneficial effects of rTPJ stimulation were demonstrated by tDCS studies (Santiesteban et al., 2012, 2015), but the evidence is mixed (Nobusako et al., 2017; Yang et al., 2020) with most of the studies applying a between-subject design lacking a baseline control.

Moreover, accuracies were substantially lower than in our study, particularly after sham or occipital stimulation, which left more room for improvement in these samples than in ours. Therefore, investigating whether clinical or subclinical samples that are characterized by decreased perspective-taking performance may benefit from neurofeedback of the rTPJ should be addressed in future studies. Lastly, more difficult perspective-taking tasks need to be designed to avoid ceiling effects in participants with high cognitive performance.

### 4.3 Secondary outcomes and non-specific mechanisms

Self-efficacy improved after the training, and, although we did not find a significant interaction effect, this effect seemed to be slightly more pronounced in the upregulation group. A number of neurofeedback studies have demonstrated improvements in domain-specific or general self-efficacy in different clinical samples and have discussed improvements of self-efficacy as a psychological mechanism mediating the effect of neurofeedback training (Schmidt and Martin, 2016, 2020; Ko and Park, 2018; Mehler et al., 2018; Hershaw et al., 2020; Markiewicz et al., 2021). We were unable to find significant correlations between changes in self-efficacy and behavioral improvements in cognitive tasks. Self-efficacy might therefore be a psychological mechanism that mediates the effects on symptom improvement in clinical samples, but this was not observed in the current sample and thus cannot be responsible for the cognitive improvement observed in the reorienting of attention task in young and healthy participants.

Regarding non-specific mechanisms, we were unable to find between-group differences in expectation towards the neurofeedback training and with respect to the evaluation of the training. Motivation slightly decreased in the upregulation group, although it remained at a high level. The decrease in motivation might be explained by the lower level of difficulty of the upregulation training compared to the downregulation training, which was experienced to be more challenging. Participants in the downregulation group also showed lower control beliefs than the upregulation group at the beginning of the training, but this difference disappeared over the course of the training once participants in the downregulation group were regulating more successfully. At the end of the training, we found a significant group difference in the amount of the monetary rewards received, which occurred due to the higher regulation success in the upregulation group. Unfortunately, subjective reward experience was not assessed in this study. It is possible, however, that higher reward experience in the upregulation group may have contributed to the differential behavioral effects in the reorienting of attention task. Indeed, we found a small, albeit insignificant, correlation of reward with improvements in this task. It is worth noting, however, that such a difference in reward experience should have affected the perspective-taking task as well, and differences in mood states, motivation, evaluation of the training, or control beliefs were not found.

In summary, these findings indicate a low influence of non-specific psychological mechanisms such as reward, treatment expectations, motivation, and control beliefs (Ros et al., 2020) and further support a neurophysiologically specific mechanism of rTPJ regulation on stimulus-driven attention.

### 4.4 Predicting neurofeedback success

The finding that improvements in stimulus-driven attention were predicted by lower baseline performance is promising for clinical translation. Clinical populations characterized by difficulties in stimulus-driven attention, for example ASD (Landry and Parker, 2013; Kana et al., 2014), may benefit more from the training than our healthy sample.

In particular, while these findings are promising from a clinical translation perspective, we acknowledge that conclusions are limited to a healthy population. Nevertheless, these exploratory findings allow us to hypothesize that measures of empathy, as well as the baseline task performance of the outcome measures, have a predictive value for the behavioral effects of neurofeedback training of the rTPJ. In this context, it is also noteworthy that comorbid impulsivity symptoms may moderate the effects of a neurofeedback intervention in ASD and should therefore also be assessed in future studies (Prillinger et al., 2022). Testing these hypotheses in confirmatory study designs including clinical samples will allow to identify and select responders of a neurofeedback intervention, which is important when it comes to the clinical translation of personalized TPJ neurofeedback protocols.

### 4.5 Limitations and future directions

This study has some limitations that are worth discussing. Since this was the first study investigating the efficacy of fNIRS neurofeedback of the rTPJ, potential effect sizes were unknown and therefore the sample size was not determined based on an a-priori statistical power analysis. While this may have resulted in insufficient statistical power to detect small effect sizes such as the hypothesized group x time interaction effect in regulation performance, it is worth underlining the large sample size in our study compared to the current state of the fNIRS neurofeedback field (Kohl et al., 2020). Secondly, both groups showed high activation from the beginning of the training and no learning effect in the upregulation group was observed, which may lead to the conclusion that mere mental rehearsal and stimulation through social stimuli is sufficient, and that neurofeedback is not necessary to regulate rTPJ activity.

In subsequent studies, it could prove advantageous to implement longer training regimes to possibly foster a learning effect in the upregulation group as well. Additionally, employing more neutral stimuli, like thermometer images, and refraining from suggesting example strategies, as well as implementing controls for the activation of a coincident neurofeedback control network, could assist in isolating a specific mechanism of rTPJ upregulation. Finally, the inclusion of extra control groups, such as a mental rehearsal group or a sham feedback group, would aid in affirming a specific mechanism as observed in the attention task.

Thirdly, this study did not involve short-distance measurement or other recommended measures of systemic physiology (Yücel et al., 2021). With the increasing availability of state-of-the art online artifact control measures and signal processing methods (Klein and Kranczioch, 2019; Lühmann et al., 2020; Klein et al., 2022; Schroeder et al., 2023), as well as hardware featuring an expanded channel count, spatial coverage and the inclusion of short-distance channels, future studies will be able to better control for systemic physiology but also for signals from irrelevant brain regions (spatial specificity). However, we took care to keep the contribution of systemic physiological changes in our design to a minimum and assessed the robustness of the online analysis through an additional offline analysis using more stringent preprocessing methods

Furthermore, care should be taken when setting and adapting feedback thresholds, particularly when differences in target regulation difficulty can be expected. Feedback thresholds were based on online assessments of rTPJ activity during the tasks before the training. We found a large variation in the assessments (range: 0.03 – 6.92), which may have been the result of suboptimal online processing methods and artifact control and may have made the training too easy or too difficult for some of the participants. Future studies can use better online processing methods and thus exclude extreme values that are likely the result of noisy measures. To avoid differences in rewards, the thresholds should be adapted more carefully throughout the training and take into account differences in regulation difficulty, as present in our bidirectional control group design, for example the downregulation group should start with lower and smaller increases of feedback thresholds.

In addition, we had some issues with blinding the participants. Participants were informed about the bidirectional control group approach and the debriefing revealed that some participants associated “downregulation” with being more difficult or less successful in the training, while “upregulation” was associated with the opposite. This might explain why more than 80% in the upregulation group guessed the group assignment correctly, although none of the participants were confident about their judgement. Notably, this lack of blinding did not seem to have an influence on participants, as evidenced by the absence of group differences in motivation, control beliefs, expectation towards the training, and evaluation of the training. Future neurofeedback experiments employing a bidirectional control group approach should take care to avoid such associations when designing the instructions for the participants. If ethically justifiable, participants should not be informed about the existence of a control condition, or at least not be informed about a downregulation condition, but rather be instructed that there are two groups in which different patterns of brain activity are reinforced.

In summary, this is the first study that demonstrated the feasibility and effectiveness of fNIRS-based neurofeedback training of the rTPJ. We present preliminary causal evidence that regulation of rTPJ activity affects stimulus-driven attention. However, it remains unclear if fNIRS-based neurofeedback can modulate social cognition. Future studies including longer training regimes and better controls are required to corroborate these initial findings in larger samples using state-of-the-art fNIRS methods. This study sets the ground for future investigations in clinical populations that are characterized by the aberrant functioning of the rTPJ or difficulties in stimulus-driven spatial attention.

## Acknowledgments

This work was supported by the German Research Foundation’s (DFG) International Research Training Group “The Neuroscience of Modulating Aggression and Impulsivity in Psychopathology” (IRTG-2150). SHK was supported by a fellowship from the Japan Society for the Promotion of Science (JSPS). The purchase of the Hitachi fNIRS system for the University Hospital RWTH Aachen (Germany) was supported by funding from the German Research Foundation (DFG; INST 948/18-1 FUGG), awarded to KK. KK is further supported by the German Research Foundation –DFG: Project-ID 431549029 –SFB 1451. DMAM is supported by a Junior Principal Investigator (JPI) fellowship funded by the Excellence Strategy of the Federal 621 Government and the Laender (grant reference number: JPI074-21). SRS is supported by European Research Council (ERC) under the project NGBMI (759370) and TIMS (101081905), the Federal Ministry of Research and Education (BMBF) under the projects SSMART (01DR21025A), NEO (13GW0483C), QHMI (03ZU1110DD) and QSHIFT (01UX2211), as well as the Einstein Foundation Berlin (A-2019-558).

We would like to thank Dr. Iroise Dumontheil for providing the stimuli of the perspective-taking task and Dr. Simone Vossel for providing the stimuli of the reorienting of attention task. We would also like to thank Mert Asil Türeli for helping to program the experimental paradigms and Růžena Ceralová for her contribution to data management.

