## Supplementary Material for "Successful Modulation of Temporoparietal Junction Activity and Stimulus-Driven Attention by fNIRS-based Neurofeedback – a Randomized Controlled Proof-of-Concept Study"

This document provides supplementary material pertaining to the manuscript “Successful Modulation of Temporoparietal Junction Activity and Stimulus-Driven Attention by fNIRS-based Neurofeedback – a Randomized, Controlled Proof-of-Concept Study”.

#### 1 Neurofeedback instructions

See Supplementary file 2 instructions.pdf

#### 2 CRED-nf checklist summary

See Supplementary file 3 CRED-nf checklist summary.pdf

**Table S1. Questionnaire results and sample characteristics**

| | **Upregulation** (*M ± SD*) | **Downregulation** (*M ± SD*) | *p-value* |
|---|---|---|---|
| **PRE** | | | |
| N | 27 (13 female) | 18 (9 female) | |
| Age (years) | 24.22 ± 3.03 | 24.22 ± 2.71 | 0.935 |
| pre rTPJ thresholds | 2.19 ± 1.45 | 2.76 ± 1.84 | 0.270 |
| AQ total | 15.30 ± 6.14 | 13.94 ± 4.49 | 0.399 |
| EQ total | 45.19 ± 9.76 | 45.17 ± 8.28 | 0.995 |
| SQ total | 29.96 ± 10.35 | 32.11 ± 13.75 | 0.577 |
| ASRS: total | 24.00 ± 9.57 | 19.94 ± 8.19 | 0.316 |
| ASRS: inattention | 12.22 ± 5.58 | 10.33 ± 4.86 | 0.236 |
| ASRS: Hyp/Imp | 11.78 ± 6.00 | 9.61 ± 4.46 | 0.172 |
| IRI total[1] | 56.78 ± 11.78 | 52.39 ± 8.83 | 0.161 |
| IRI perspective-taking | 20.07 ± 3.92 | 18.06 ± 2.94 | 0.055 |
| IRI fantasy | 16.78 ± 5.20 | 16.11 ± 5.12 | 0.673 |
| IRI empathic concern | 19.93 ± 4.51 | 18.22 ± 5.00 | 0.252 |
| IRI personal distress | 10.89 ± 2.94 | 9.61 ± 4.94 | 0.333 |
| SRS: total | 39.26 ± 19.36 | 37.94 ± 12.94 | 0.862 |
| SRS: social awareness | 5.00 ± 2.73 | 4.89 ± 2.03 | 0.921 |
| SRS: social cognition | 7.00 ± 3.99 | 6.06 ± 3.42 | 0.401 |
| SRS: social communication | 11.85 ± 9.07 | 10.83 ± 4.54 | 0.871 |
| SRS: social motivation | 7.89 ± 4.23 | 7.50 ± 3.54 | 1.000 |
| SRS: autistic mannerism | 7.11 ± 4.12 | 8.06 ± 3.59 | 0.334 |
| Expectations | 2.42 ± 0.56 | 2.67 ± 0.59 | 0.123 |
| Motivation | 3.70 ± 0.40 | 3.63 ± 0.50 | 0.695 |
| **POST** | | | |
| Evaluation: belief efficacy | 2.40 ± 0.87 | 2.39 ± 0.83 | 0.861 |
| Evaluation: joy | 2.13 ± 0.35 | 2.22 ± 0.40 | 0.431 |
| Evaluation: experimenter | 3.87 ± 0.26 | 3.81 ± 0.35 | 0.610 |
| Neurofeedback control belief | 7.59 ± 0.93 | 6.56 ± 1.54 | 0.020 |
| Monetary reward | 12.80 ± 2.33 | 7.86 ± 2.16€ | < 0.001 |

[1] according to Cliffordson (2001) and Paulus (2012); AQ, Autism Spectrum Quotient; ASRS, Adult ADHD Self-Report Scale; EQ, Empathy Quotient; IRI, Interpersonal Reactivity Index; POMS, Profile of Mood States; SQ, Systemizing Quotient; SRS, Social Responsiveness Scale

#### Table S2. Descriptive statistics of pre-post experimental tasks

| | Pre *(M ± SD)* | Post *(M ± SD)* |
|---|---|---|
| **Upregulation group** | | |
| RTs attention task – invalid | 497 ± 69 ms | 481 ± 57 ms |
| RTs attention task – valid | 452 ± 61 ms | 433 ± 47 ms |
| Accuracies attention task – invalid | 0.98 ± 0.02 | 0.98 ± 0.03 |
| Accuracies attention task – valid trials | 0.99 ± 0.02 | 0.99 ± 0.03 |
| RTs vPT task – PT | 3667 ± 327 ms | 3520 ± 282 ms |
| RTs vPT task – NPT | 3670 ± 286 ms | 3490 ± 296 ms |
| Accuracies vPT task – PT | 0.94 ± 0.07 | 0.98 ± 0.04 |
| Accuracies vPT task – NPT | 0.98 ± 0.04 | 0.99 ± 0.03 |
| **Downregulation group** | | |
| RTs attention task – invalid | 508 ± 90 ms | 521 ± 110 ms |
| RTs attention task – valid | 468 ± 94 ms | 486 ± 106 ms |
| Accuracies attention task – invalid | 0.98 ± 0.05 | 0.98 ± 0.05 |
| Accuracies attention task – valid trials | 0.99 ± 0.03 | 0.99 ± 0.02 |
| RTs vPT task – PT | 3630 ± 302 ms | 3500 ± 280 ms |
| RTs vPT task – NPT | 3620 ± 272 ms | 3480 ± 232 ms |
| Accuracies vPT task – PT | 0.93 ± 0.09 | 0.98 ± 0.03 |
| Accuracies vPT task – NPT | 0.97 ± 0.04 | 0.99 ± 0.02 |

Note that we did not find a significant a three-way interaction of group × time × condition in the attention task or the vPT and only a group × time interaction in the attention task. Reaction times across conditions decreased in the upregulation group (pre = 474 ± 68ms, post = 457 ± 57ms, d = 0.51) and increased in the downregulation group across conditions (pre = 488 ± 93ms, post = 503 ± 108ms, d = -0.56; see main text). PT, perspective taking; NPT, non-perspective taking; vPT, visual perspective taking.

#### Table S3. POMS and general self-efficacy

| | Pre | Post | p-value |
|---|---|---|---|
| | *M ± SD or Md (IQR)* | | |
| **Upregulation group** | | | |
| General self-efficacy | 30.59 ± 2.50 | 32.04 ± 3.16 | 0.068 |
| POMS: depression/anxiety | 0.21 (0.54) | 0.14 (0.43) | 0.104 |
| POMS: vigor | 3.43 (1.14) | 3.29 (0.86) | 0.592 |
| POMS: fatigue | 1.61 ± 0.99 | 1.81 ± 1.20 | 0.492 |
| POMS: hostility | 0.14 (0.21) | 0.14 (0.21) | 0.439 |
| **Downregulation group** | | | |
| General self-efficacy | 31.83 ± 3.57 | 32.00 ± 3.41 | 0.887 |
| POMS: depression/anxiety | 0.11 (0.63) | 0.18 (0.29) | 0.277 |
| POMS: vigor | 3.40 ± 0.90 | 2.94 ± 1.31 | 0.229 |
| POMS: fatigue | 1.86 (1.71) | 1.57 (1.86) | 0.228 |
| POMS: hostility | 0.07 (0.93) | 0.21 (0.79) | 0.627 |

#### Table S4. Motivation and self-control beliefs throughout the training

| | Session 1 | Session 2 | Session 3 | Session 4 | *F* | *p* |
|---|---|---|---|---|---|---|
| | *M ± SD or Md (IQR)* | | | | | |
| **Upregulation group** | | | | | | |
| Motivation | 9 (2.25) | 9 (2) | 9 (2.25) | 8 (3) | 2.54 | 0.021* |
| Self-control belief | 7.19±1.42 | 6.92±1.5 | 7.08±1.35 | 6.88±1.82 | 0.11 | 0.936 |
| **Downregulation group** | | | | | | |
| Motivation | 9.5 (1.75) | 9 (1.75) | 9 (2) | 9.5 (1.75) | 0.681 | 0.48 |
| Self-control belief | 4.44±1.82 | 5.72±1.64 | 6.56±1.82 | 6.36±2.11 | 4.75 | 0.005** |

#### 3 Neurofeedback regulation success

##### 3.1 Regulation behavior – durations above feedback thresholds

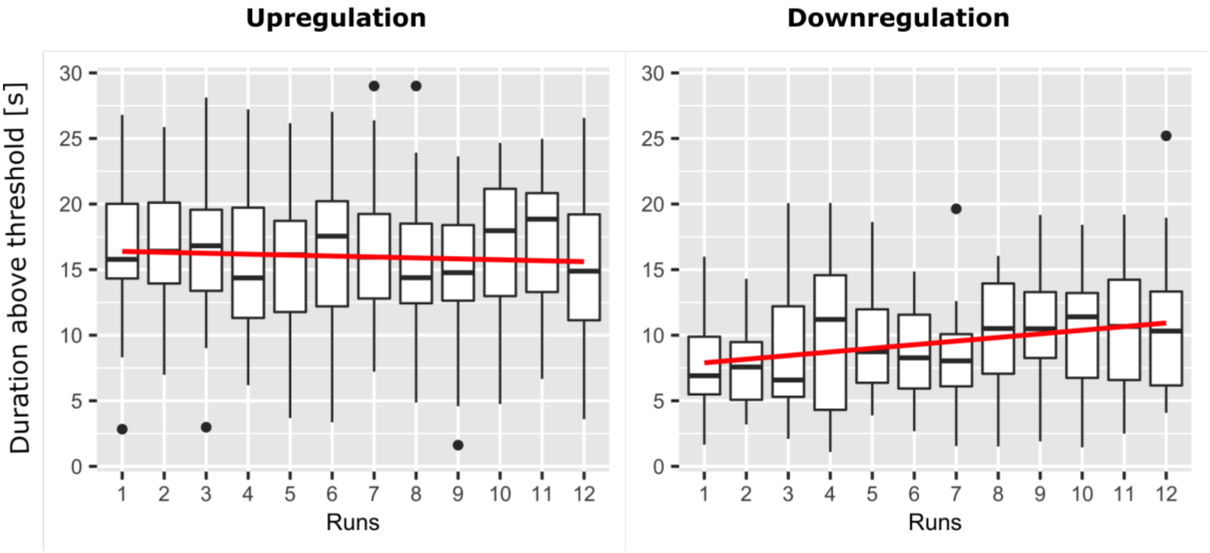


**Figure S1. Development of feedback performance over time (maximum durations above the threshold).** Box plots depicting the feedback performance as assessed by the maximum durations that participants could sustain the feedback signal above the individual threshold averaged over all participants for each run. The regression lines of the linear models are depicted in red.

Figure S1 shows the development of feedback performance over time (maximum durations above the threshold in a trial averaged over all participants for each run). Participants were able to keep the feedback signal above their individual thresholds and we found similar results to the main analysis of regulation success (analysis based on signal amplitudes). One-sample t-tests revealed a significant effect of regulation in the upregulation group ($M$ = 16±3.25s, t(26) = 25.59, $p < 0.001$, $d = 5.02$) and downregulation group ($M$ = 9.41±2.9s, $t(17) = 13.76$, $p < 0.001$, $d = 3.34$). Paired-sample t-tests only revealed a difference between the last and the first session in the downregulation group ($Mdiff$ = 3.2±4.78, $t(17) = 2.83$, $p$ = 0.01, $d = 0.56$). This was not the case in the upregulation group ($Mdiff$ = -0.86±5.89s, $p$ = 0.46, $d = 0.15$). Mixed models revealed a significant effect of groups ($F(1,43) = 48.25$, $p <$ 0.001, $\eta_p^2 = 0.53$) as well as a marginal group × time effect ($F(11,473) = 1.72$, $p = 0.07$, $\eta_p^2$ = 0.04) and time effect ($F(11,473) = 1.53$, $p = 0.12$, $\eta_p^2 = 0.03$). There was no significant within-group time effect in either group. In the upregulation group, 40.7% of the participants (11 of 27) showed increasing durations above the threshold over trials, as indicated by a positive slope of the regression, and 16 of 27 participants (59.26%) showed longer durations above the threshold in the last session compared to the first session. In the downregulation group, 61.1% of the participants (11 of 18) showed increasing durations above the threshold over trials, as indicated by a positive slope of the regression and twelve of 18 participants (66.7%) showed longer durations above the threshold in the last session compared to the first session.

##### 3.2 Dynamic of the feedback signal – switching between above and below the threshold

We also explored the dynamics of the feedback signal by analyzing the number of switches between below and above the feedback threshold, but observed no significant effects (see Figure S2). Mixed models revealed no significant interaction effect or main effect of time. On average, participants switched 1.92 times per trial (upregulation group = 2 ± 0.10, downregulation group = 1.80 ± 0.18 times). Paired-sample t-tests comparing the last and the first session also revealed no effect for either group. In the upregulation group, 48.15% of the participants (13 of 27) showed decreasing number of switches between below and above the threshold over trials, as indicated by a negative slope of the regression, and 51.85% (14 of 27 participants) showed less switching in the last session compared to the first session. In the downregulation group, 55.56% of the participants (10 of 18) showed decreasing switches over trials, as indicated by a negative slope of the regression, and 38.89% (7 of 18 participants) showed less switching in the last session compared to the first session.

In total, the number of switches between above and below a feedback threshold did not reveal a learning effect. On average, participants did not cross the feedback threshold very often (only 1-2 times per trial), which made this measure unsuitable for detecting a learning effect. However, the sensitivity of time-based measures depends on the appropriate estimation and selection of feedback thresholds used for the calculation (see limitations).

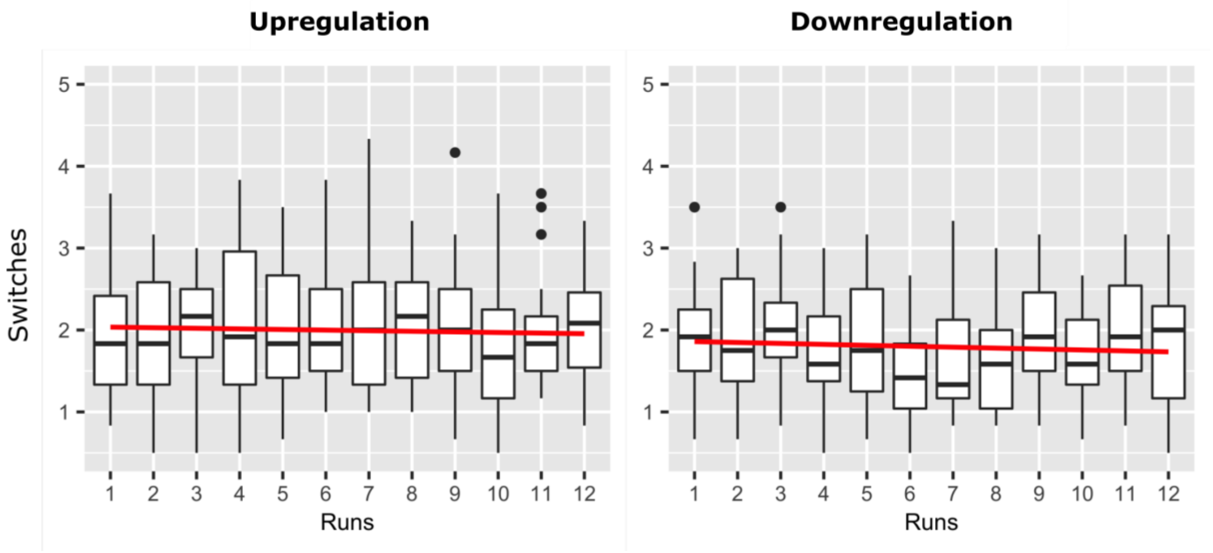


**Figure S2. Dynamic of the feedback signal.** Box plots depicting the number of switches between above and below the threshold averaged over all participants for each run. The regression lines of the linear models are depicted in red.

#### Table S5. Individual neurofeedback success in the upregulation group

| Subject | Succesful runs | | Last vs. first | | Linear slope | |
|---|---|---|---|---|---|---|
| U1 | 7 | Yes | -2.28 | No | -0.09 | No |
| U2 | 12 | Yes | 3.00 | Yes | 0.21 | Yes |
| U3 | 10 | Yes | -3.51 | No | -0.32 | No |
| U4 | 8 | Yes | 0.70 | Yes | 0.00 | Yes |
| U5 | 10 | Yes | 3.91 | Yes | 0.13 | Yes |
| U6 | 9 | Yes | 5.16 | Yes | 0.62 | Yes |
| U7 | 8 | Yes | 0.79 | Yes | -0.13 | No |
| U8 | 11 | Yes | -3.84 | No | -0.36 | No |
| U9 | 7 | Yes | -7.18 | No | -0.59 | No |
| U10 | 12 | Yes | 0.76 | Yes | 0.30 | Yes |
| U11 | 12 | Yes | 2.56 | Yes | 0.17 | Yes |
| U12 | 12 | Yes | 4.46 | Yes | 0.43 | Yes |
| U13 | 12 | Yes | 2.39 | Yes | 0.16 | Yes |
| U14 | 10 | Yes | 0.63 | Yes | 0.00 | No |
| U15 | 10 | Yes | -4.92 | No | -0.41 | No |
| U16 | 5 | No | 7.75 | Yes | 0.81 | Yes |
| U17 | 8 | Yes | -1.75 | No | -0.13 | No |
| U18 | 11 | Yes | -1.70 | No | 0.03 | Yes |
| U19 | 11 | Yes | -6.86 | No | -0.34 | No |
| U20 | 12 | Yes | 3.93 | Yes | 0.16 | Yes |
| U21 | 12 | Yes | 2.10 | Yes | 0.16 | Yes |
| U22 | 7 | Yes | 1.54 | Yes | 0.00 | Yes |
| U23 | 12 | Yes | -3.00 | No | -0.26 | No |
| U24 | 7 | Yes | -0.17 | No | -0.25 | No |
| U25 | 7 | Yes | 1.75 | Yes | 0.45 | Yes |
| U26 | 6 | Yes | -3.43 | No | -0.44 | No |
| U27 | 12 | Yes | -2.14 | No | -0.10 | No |
| | M = 9.63 | 26/27 | M = 0.02 ± 3.7 | 15/27 | M = 0.01 ± 0.34 | 14/27 |

**Table S6. Individual neurofeedback success in the downregulation group**

| Subject | Succesful runs | | Last vs. first | | Linear slope | |
|---|---|---|---|---|---|---|
| D1 | 2 | No | -2.18 | Yes | -0.04 | Yes |
| D2 | 6 | Yes | -2.93 | Yes | -0.04 | Yes |
| D3 | 6 | Yes | -8.68 | Yes | -0.20 | Yes |
| D4 | 2 | No | -0.14 | Yes | -0.03 | Yes |
| D5 | 0 | No | -2.03 | Yes | -0.06 | Yes |
| D6 | 0 | No | -0.24 | Yes | -0.01 | Yes |
| D7 | 5 | No | -1.51 | Yes | 0.02 | No |
| D8 | 0 | No | -1.05 | Yes | -0.03 | Yes |
| D9 | 9 | Yes | -8.99 | Yes | -0.09 | Yes |
| D10 | 4 | No | -1.28 | Yes | -0.01 | Yes |
| D11 | 3 | No | -8.40 | Yes | -0.12 | Yes |
| D12 | 3 | No | -2.35 | Yes | -0.03 | Yes |
| D13 | 0 | No | -4.32 | Yes | -0.06 | Yes |
| D14 | 0 | No | -0.91 | Yes | -0.03 | Yes |
| D15 | 3 | No | 0.51 | No | -0.03 | Yes |
| D16 | 6 | Yes | 0.34 | No | 0.01 | No |
| D17 | 4 | No | -3.91 | Yes | -0.04 | Yes |
| D18 | 1 | No | 5.55 | No | 0.08 | No |
| | M = 3 | 4/18 | M = -2.36 ± 3.6 | 15/18 | M = -0.22 ± 0.35 | 15/18 |

#### 3.3 Robustness checks

Figure S3 shows the regulation performance based on the robustness checks, Table S7 statistical results and figure S4 the grand averages based on the different analyses approaches.

##### Robustness check 1: Bandpass filter (0.01-0.09Hz)

Robustness check 1 confirmed the results of the online analysis. One-sample t-tests again revealed a significant effect of regulation in the upregulation group (*M* = 5.02±4.74, t(26) = 5.51, $p < 0.001$, *d* = 1.08) and downregulation group (*M* = 2.39±4.28, $t(17) = 2.37$, $p = 0.03$, d = 0.58). Paired-sample t-tests only revealed a difference between the last and first session in the downregulation group (*Mdiff* = -6.22±7.76, $t(17) = 2.79$, $p = 0.01$, *d* = 0.82). This was not the case in the upregulation group (*Mdiff* = -1.03±8.59, $p = 0.54$, *d* = 0.12). The non-parametric ANOVA revealed a marginal group effect ($F_{ATS}$ (1, ∞) = 3.33, $p = 0.07$), no significant group × time interaction, and a significant time effect ($F_{ATS}$ (8.17, ∞) = 2.23, *p* =

0.02). Separate analysis for both groups showed no effect in the upregulation group and only a non-significant time trend in the downregulation group ($F$(11,204) = 1.59, $p$ = 0.1, $\eta_p^2$ = 0.08).

##### Robustness check 2: Common average reference

On the group level, none of the effects survived the more conservative robustness check 2. One-sample t-tests revealed no significant effect of regulation in the upregulation group ($M$ = -0.15±2.76, $p$ = 0.78, $d$ = -0.06), and no significant effect in the downregulation group ($M$ = 0.11±4.68, $p$ = 0.93, $d$ = 0.02). Paired-sample t-tests revealed no difference between the last and the first session in both groups. The learning effect for the downregulation group disappeared ($Mdiff$ = -1.25±5.28, $p$ = 0.329, $d$ = 0.24). The non-parametric ANOVA revealed no group effect and no significant group × time interaction. The trend effect of time disappeared (both groups: $F_{ATS}$ (8.4, ∞) = 0.87, $p$ = 0.52; downregulation group: $F_{ATS}$ (6.68, ∞) = 0.91, $p$ = 0.5).

##### Table S7. Neurofeedback regulation success – offline robustness checks

| | Offline robustness check 1 (BP 0.01-0.09Hz) | | | Offline robustness check 2 (CAR) | | |
|---|---|---|---|---|---|---|
| **Upregulation group** | | | | | | |
| | *M±SD* | *N* | *p-value* | *M±SD* | *N* | *p-value* |
| NF performance - compared to baseline | 5.02±4.74 | 25/27 | 0.001* | -0.15±2.76 | 15/27 | 0.612 |
| NF improvement (slope) | -0.11±0.83 | 11/27 | 0.25 | -0.02±0.45 | 12/27 | 0.43. |
| NF improvement (last vs first) | 2.39±4.28 | 13/27 | 0.54 | 0.20±5.39 | 13/27 | 0.845 |
| **Downregulation group** | | | | | | |
| | *M±SD* | *N* | *p-value* | *M±SD* | *N* | *p-value* |
| NF performance - compared to baseline | 2.39±4.28 | 7/18 | 0.99 | 0.11±4.68 | 10/18 | 0.537 |
| NF improvement (slope) | -0.40±0.76 | 14/18 | 0.1 | -0.12±0.47 | 14/18 | 0.496 |
| NF improvement (last vs first) | -6.22±7.76 | 14/18 | 0.003* | -1.25±5.28 | 11/18 | 0.329 |

Neurofeedback regulation success according to different success measures and offline robustness checks for both groups. The p-values reflect the results of the group analysis based on the description in 2.5 “Data processing and analysis”; NF, neurofeedback

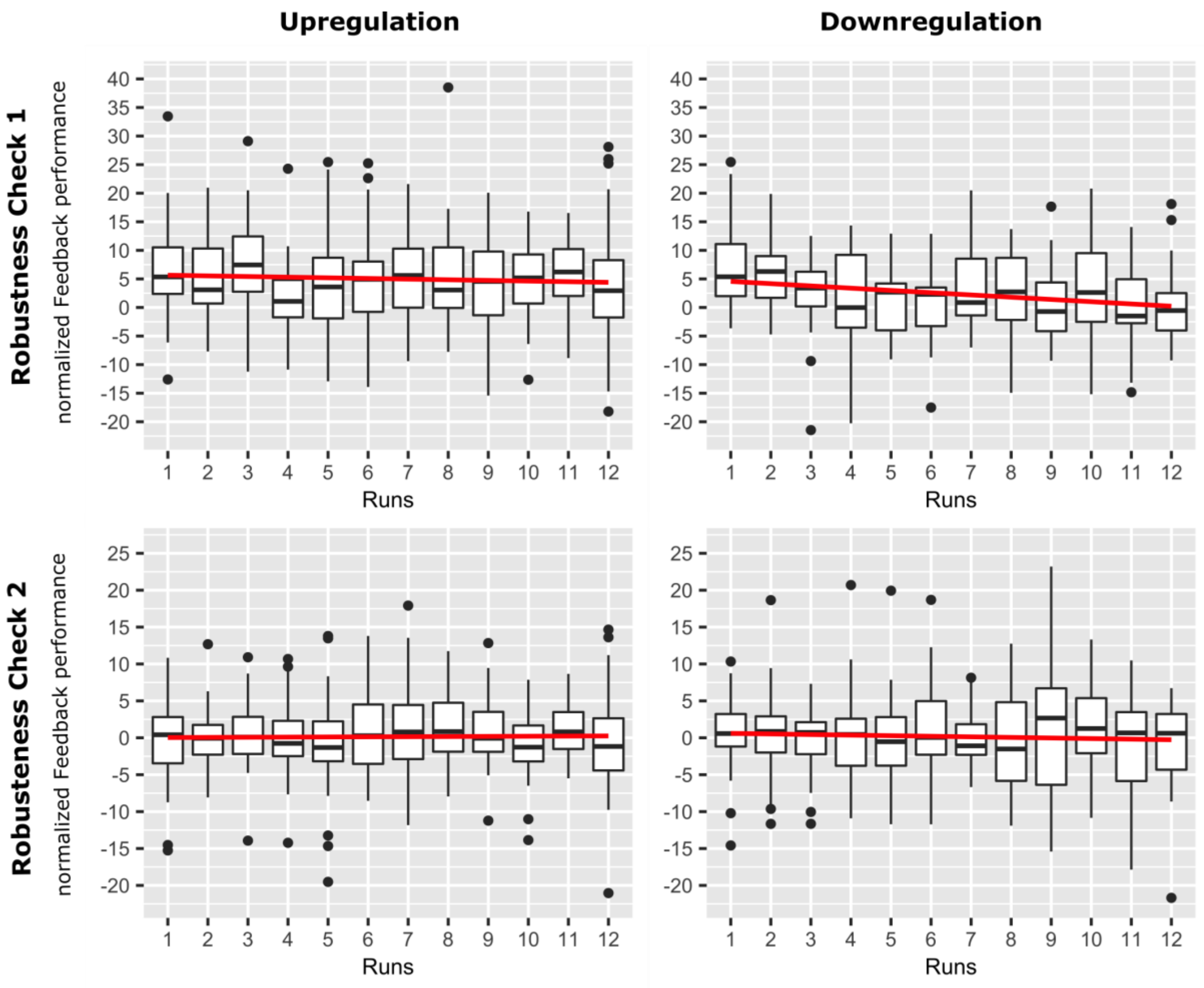


**Figure S3. Neurofeedback regulation performance based on an additional offline analysis using more stringent artifact correction methods (robustness checks).** The first row shows box plots of the average feedback performance as assessed by the standardized median change of rTPJ activation averaged over participants for all runs based on robustness check 1 (stronger bandpass filter 0.01-0.09Hz, see methods section). The second row shows box plots of the average feedback performance as assessed by the standardized median change of rTPJ activation averaged over participants for all runs based on robustness check 2 (common average correction, see methods section). The regression lines of the linear models are depicted in red.

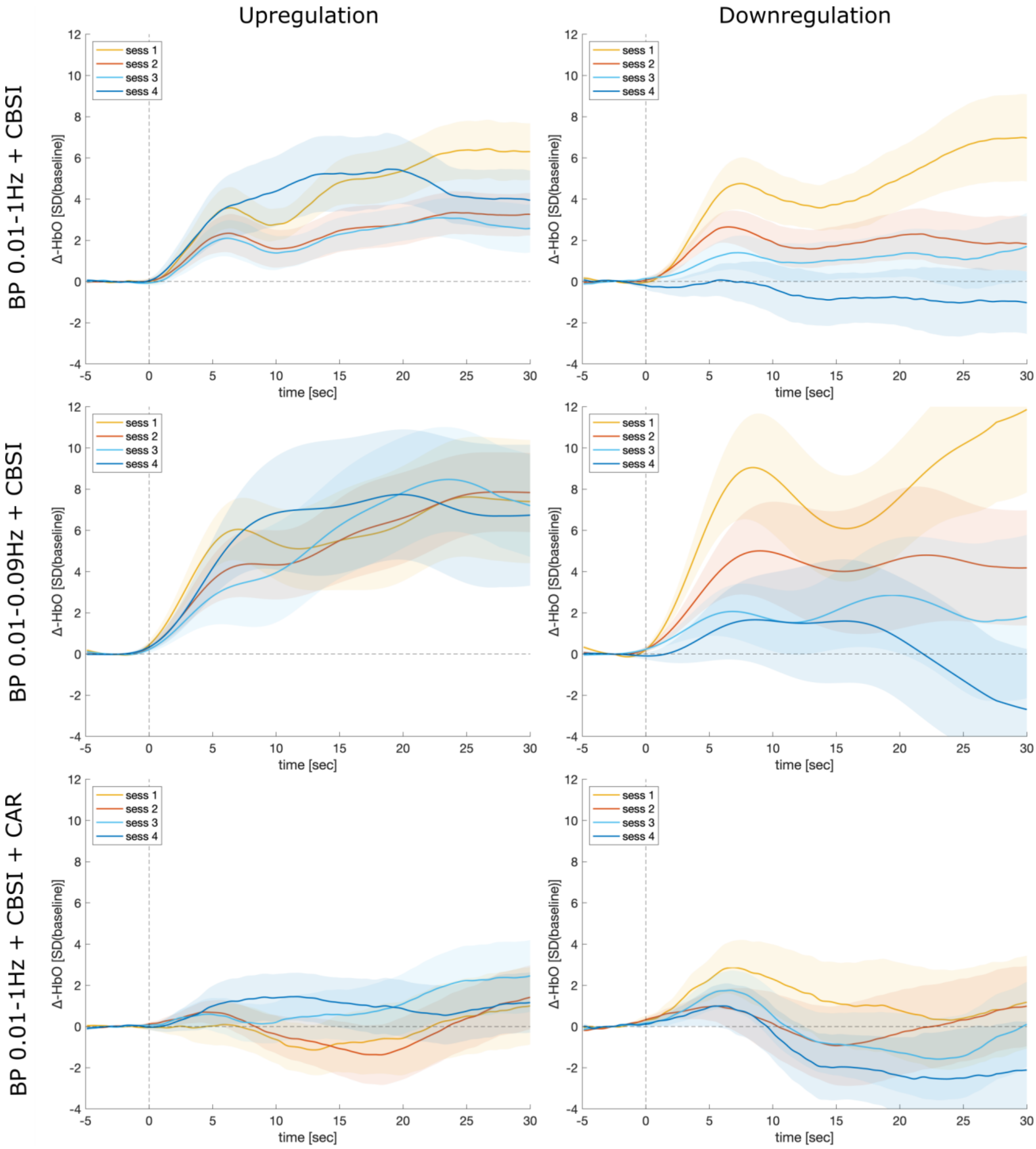


**Figure S4. Grand averages of the feedback channel for the sessions of both groups.** The three rows show the grand averages ± standard error (shaded area) for the three different offline analysis approaches based on the raw fNIRS signal. First row: offline analysis mimicking the online analysis; second row: analysis with a more stringent bandpass filter; third row: common average reference (CAR) approach. All analysis approaches included the correlation-based signal improvement (CBSI; Cui et al., 2010) and a 5s moving average filter.

#### 4 Results for the reorienting of attention task including valid only blocks

The pre-post comparisons revealed that after the neurofeedback training, reaction times decreased in the upregulation group (pre = 468 ± 71ms, post = 450 ± 61ms, $d = 0.58$) and increased in the downregulation-group across conditions (pre = 480 ± 94ms, post = 493 ± 107ms, $d = -0.49$), as indicated by a significant group × time interaction ($F(1,123) = 15.14$, $p < 0.001$) and a significant main effect of time in the upregulation group ($F_{ATS}(1, \infty) = 10.01$, $p = 0.001$) as well as a marginally significant effect of time in the downregulation group ($F_{ATS}(1, \infty) = 3.25$, $p = 0.071$). No three-way interaction of group × time × condition was observed.

#### 5 Mental strategies underlying neurofeedback regulation

Table S8 shows the results for the reported mental strategies and their success ratings for both groups relative to the total number of strategies reported by each group. In total, mental strategies for 1299 trials were reported (771 by the upregulation group and 528 by the downregulation group). Socio-cognitive strategies and positive mental imagery were reported the most in both groups. Both groups seemed to be equally reliant on socio-cognitive strategies (upregulation group: 15.82% vs. downregulation group: 15.34), with the upregulation group reporting socio-cognitive strategies as more effective than the downregulation group (mean rating: 4.03 vs. 2.28). The upregulation group used more positive mental imagery, arithmetic strategies, and music-related strategies than the downregulation group. The downregulation group used slightly more relaxation strategies and strategies related to memory and focused attention.

Most strategies seemed to work better in the upregulation group (mean success rating of 3.35) than in the downregulation group (mean success rating of 2.74). Socio-cognitive, arithmetic, and working memory strategies seemed to perform better in the upregulation group. Moreover, the upregulation group reported positive and negative mental imagery, language, music, and visual imagery as slightly more effective. The downregulation group reported thinking, memory, and relaxation strategies as slightly more effective. In summary, both groups did not differ much regarding the use of mental strategies, but some strategies seemed to work better depending on the regulation condition.

#### Table S8. Reported mental strategies and their success ratings for both groups

| | Upregulation group | | Downregulation group | |
|---|---|---|---|---|
| | Strategies reported [%] | Mean rating | Strategies reported [%] | Mean rating |
| Socio-cognitive strategies | 15.40 | 4.11 | 15.34 | 2.28 |
| Positive mental imagery | 24.05 | 3.42 | 16.86 | 2.99 |
| Language | 6.45 | 3.43 | 6.25 | 3.05 |
| Counting | 6.60 | 2.98 | 5.11 | 2.59 |
| Arithmetic | 9.24 | 3.31 | 5.68 | 2.10 |
| Working memory | 5.72 | 3.32 | 5.49 | 3.03 |
| Planning | 2.35 | 2.88 | 1.70 | 3.39 |
| Desire to regulate and increase feedback signal | 4.69 | 3.25 | 5.49 | 3.14 |
| Out-of-body imagination | 2.79 | 2.71 | 4.92 | 2.65 |
| Music | 6.60 | 3.01 | 3.98 | 2.71 |
| Relaxation | 2.79 | 3.08 | 8.33 | 3.45 |
| Negative mental imagery | 0.59 | 2.75 | 1.14 | 2.42 |
| Motor imagery | 2.35 | 2.56 | 1.70 | 2.78 |
| Connection to avatar | 1.47 | 3.05 | N/A | N/A |
| Thinking | 0.44 | 1.67 | 1.33 | 2.71 |
| Memory | 1.61 | 3.09 | 4.73 | 3.40 |
| Focused attention | 0.59 | 2.75 | 5.30 | 2.77 |
| Visual imagery | 5.13 | 3.44 | 4.73 | 2.70 |
| Sensory imagery | N/A | N/A | 0.38 | 2.50 |
| Mindfulness | N/A | N/A | 0.95 | 4.20 |
| Uncategorized | 1.17 | 2.50 | 0.57 | 3.67 |
| **Total** | | 3.35 | | 2.84 |

### 6 Correlations of behavioral outcomes with regulation performance and psychosocial factors

#### Table S9. Significant correlations between behavioral outcomes and regulation success measures with and without multiple comparison correction

| **Group** | **Behavioral outcome** | **Regulation success measure** | **Rho** | **p uncorrected** | **p adjusted** |
|---|---|---|---|---|---|
| both | **Δ** RT attention | successful runs | -0.38 | 0.012 | 0.321 |
| both | **Δ** RT attention - valid | succesful runs | -0.47 | 0.002 | 0.045* |
| both | **Δ** accuracy perspective taking | slopes | -0.38 | 0.012 | 0.327 |
| both | **Δ** accuracy perspective taking - NPT | slopes | -0.49 | 0.001 | 0.021* |
| up | **Δ** RT attention | slopes | 0.41 | 0.041 | 1.000 |
| up | **Δ** RT attention - invalid | slopes | 0.39 | 0.047 | 1.000 |
| up | **Δ** accuracy perspective taking | successful runs | -0.39 | 0.045 | 1.000 |

| | | | | | |
|---|---|---|---|---|---|
| up | **Δ** accuracy perspective taking - PT | successful runs | -0.42 | 0.030 | 0.799 |
| down | **Δ** accuracy perspective taking - NPT | slopes | -0.71 | 0.001 | 0.039* |

* significant after Bonferroni–Holm correction; NPT, non-perspective taking trials; PT, perspective taking trials.

#### Table S10. Significant correlations between behavioral outcomes and psychosocial factors with and without multiple comparison correction

| Group | Behavioral outcome | Psychosocial factor | rho | p uncorrected | p adjusted |
|---|---|---|---|---|---|
| both | **Δ** RT attention | NF control belief | -0.3 | 0.048 | 1.000 |
| both | **Δ** RT attention | Monetary reward | -0.36 | 0.020 | 0.650 |
| both | **Δ** accuracy perspective taking | Expectations | 0.33 | 0.027 | 0.894 |
| up | **Δ** RT attention | NF control belief | -0.5 | 0.041 | 1.000 |
| down | **Δ** RT attention | Evaluation: experimenter | -0.46 | 0.019 | 0.637 |

* significant after Bonferroni–Holm correction; NF, neurofeedback
