## Supplementary Material 1 for "Successful Modulation of Temporoparietal Junction Activity and Stimulus-Driven Attention by fNIRS-based Neurofeedback – a Randomized Controlled Proof-of-Concept Study"

### Neurofeedback instructions (translated from German)

During this task you will receive Feedback of the activation of a certain brain region with the goal for you to learn how to control it. The feedback will be delivered in the form of a smiling avatar. The better you are at regulating, the more the avatar will smile.

One neurofeedback run consists of 6 block. There will be 2 to 4 runs during each session.

A block consists of a regulation and a non-regulation condition.

During the non-regulation condition, you will see the avatar with a neutral facial expression. You will not receive any feedback, so the avatar will not smile. Next to the avatar you will see two + signs, which indicate the condition. Please, try not to regulate your brain activity during this condition. Just look at the avatar, relax and don’t think about anything specific.

After that the non-regulation condition will start, which you will notice through two 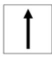 appearing next to the avatar. Try now to regulate your brain activity by using mental strategies and make the avatar smile.

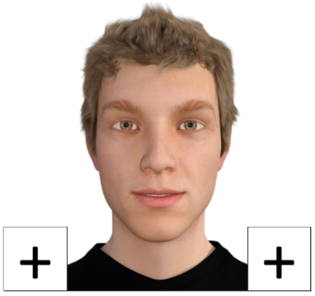

Nicht regulieren
und entspannen (20-25 sec)

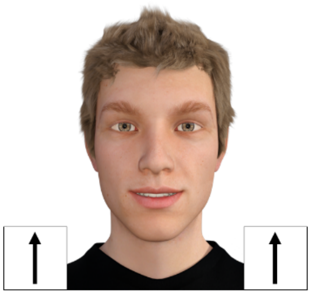

Regulierung (30 sec)

Depending on how good you are at regulating your brain activity you can earn additional money.

As soon as your brain activity passes a certain threshold a green frame will appear around the avatar. For each second that you are able to remain above this threshold you will earn 0.01€. At the end of each block you will see how much you have earned.

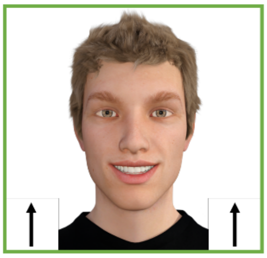

***Strategies***

As already said, you should use mental strategies to regulate your brain activity.

It is important that you find your own individual strategy to succeed. These can be different for everyone. Here are a few examples that you could try to start with:

- Try to put yourself into the avatar. What is he currently thinking or feeling?
- Think about what he is doing in his life. For example, what is his job? What is he studying? What kind of hobbies does he have? What did he do yesterday? How was his day?
- Imagine you can make the avatar smile with your own mental states
- Think about you being the avatar
- Think about leaving your body
- Think about positive life events
- Imagine a simple arithmetic problem
- Count backwards in steps of 7.
- Imagine words and read them backwards

If these do not work for you, you can also come up with your own strategies and try them out. Anything that helps to make the avatar smile is good.

Note that the signal is delayed about 4-6 seconds. So use only one strategy during one block and don’t switch in between.

Again, please only use mental strategies. Stay calm and relaxed and breath regularly.

Good Luck!

### Neurofeedback instruction (in German)

In dieser Aufgabe erhältst du Rückmeldung über die Gehirnaktivierung eines bestimmten Areals und sollst lernen diese zu beeinflussen. Die Rückmeldung wird dir in From eines lächelnden Avatars gegeben. Je besser du regulierst, desto mehr lächelt der Avatar.

Ein Neurofeedback-Durchgang besteht aus 6 Blöcken. Insgesamt besteht das Neurofeedback aus 2-4 Durchgängen.

Ein Block besteht jeweils aus einer Nichtregulierungs- und Regulierungsbedingung.

Während der Nichtregulierungsbedingung siehst du den Avatar mit neutralem Gesichtsausdruck. Du erhältst in dieser Bedingung kein Feedback, der Avatar wird also nicht lächeln. Neben dem Avatar siehst du zwei + Zeichen, die dir die Bedingung anzeigen. Bitte versuche in dieser Bedingung nicht deine Gehirnaktivität zu regulieren. Schaue einfach den Avatar an, entspanne dich und denke an nichts bestimmtes.

Danach beginnt die Regulierungsbedingung. Das erkennst du daran, dass zwei 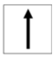 neben dem Avatar erscheinen. Versuche jetzt mittels mentaler Strategien, deine Hirnaktivierung zu regulieren und den Avatar zum Lächeln zu bringen.

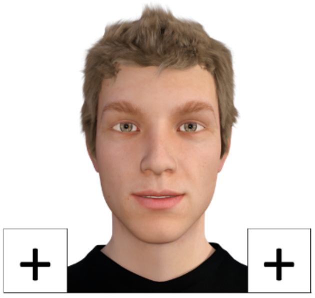

Nicht regulieren
und entspannen (20-25 sec)

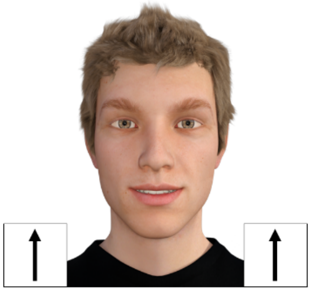

Regulierung (30 sec)

Je nachdem wie gut du regulieren kannst, kannst du in dieser Aufgabe Geld dazugewinnen.

Sobald deine Hirnaktivierung eine bestimmte Schwelle überschritten hat erscheint ein grüner Rahmen um den Avatar. Für jede Sekunde, die es dir gelingt, über dieser Schwelle zu bleiben, erhälst du 0,01 €. Am Ende eines jden Blocks wir dir dann angezeigt, wieviel du gewonnen hast.

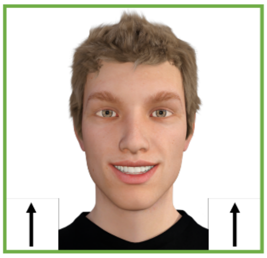

***Strategien***

Wie schon gesagt, sollst du mentale Strategien nutzen, um deine Hirnaktivierung zu regulieren.

Wichtig ist, dass du deine eigene für dich erfolgreiche Strategie findest. Das kann bei jedem eine andere sein. Hier einige Beispiele, die du am Anfang ausprobieren kannst:

- Sich in den Avatar hineinversetzen. Zum Beispiel überlegen, was dieser gerade denkt oder wie er sich fühlt.
- Überlegen, was der Avatar macht. Z.B was macht er beruflich? Was studiert er? Was hat er für Hobbies? Was hat er gestern gemacht, wie war sein Tag?
- Die Vorstellung, den Avatar mit Hilfe der eigenen mentalen Zustände zum lächeln zu bringen.
- Vorstellung, dass man selber der Avatar ist.
- Vorstellung, dass man seinen Körper verlässt.
- Sich positive eigene Lebensereignisse vorstellen.
- Einfache Rechenaufgabe vorstellen.
- Zählen, z.B. in 7er Schritten Rückwarts zählen.
- Wörter vorstellen, die man rückwärts liest.

Falls diese nicht funktionieren, kannst du dir aber auch eigene Strategien überlegen und diese ausprobieren. Wichtig ist, dass der Avatar lächelt.

Das Signal ist um 4-6 Sekunden verzögert. Probiere also pro Block (30sek) nur eine Strategie aus und wechsele nicht mittendrin.

Nochmal zur Erinnerung: Nutze nur mentale Strategien. Bleibe ruhig und entspannt sitzen und atme regelmäßig.

Viel Erfolg!
