## Supplementary Material 2 for "Successful Modulation of Temporoparietal Junction Activity and Stimulus-Driven Attention by fNIRS-based Neurofeedback – a Randomized Controlled Proof-of-Concept Study"

### CRED-nf checklist summary

*11 July, 2023*

**Manuscript title:** Successful Modulation of Temporoparietal Junction Activity and Stimulus-Driven Attention by fNIRS-based Neurofeedback – a Randomized Controlled Proof-of-Concept Study
**Corresponding Author:** Simon H. Kohl

| Item No. | Checklist item | Manuscript Details |
|---|---|---|
| **Pre-experiment** | | |
| 1a | Pre-register experimental protocol and planned analyses | *This experiment was not preregistered* |
| 1b | Justify sample size | This was a proof-of-concept study. Hence, no a priori power analysis was conducted. However, according to a sensitivity analysis, a mixed analysis of variance (ANOVA) including 45 participants was sufficiently powered (80%) to detect a group x time interaction effect of at least f = 0.43 (assuming no violation of sphericity and a correlation among repeated measure of 0.8) or 0.77 for an independent t-test. |
| **Control groups** | | |
| 2a | Employ control group(s) or control condition(s) | In a bidirectional regulation control group design, 50 healthy participants were either reinforced to up- or downregulate rTPJ activation over four days of training. |
| 2b | When leveraging experimental designs where a double-blind is possible, use a double-blind | *The experiment did not include a double-blind* |
| 2c | Blind those who rate the outcomes | *Those who rated the outcome were not blind to group assignment* |
| | Blind those who analyse the data | *Those who analysed the data were not blind to group assignment* |
| 2d | Examine to what extent participants and experimenters remain blinded | Furthermore, they were asked to guess the group condition they had been randomly assigned to. |
| 2e | In clinical efficacy studies, employ a standard-of-care intervention group as a benchmark for improvement | *NA: This is not a clinical efficacy study* |

| | | |
|---|---|---|
| 3a | Collect data on psychosocial factors | At the end of the session, participants filled in the general self-efficacy scale again as well as a debriefing questionnaire to further assess feasibility and unspecific mechanisms. This questionnaire included items assessing participants' evaluation of the neurofeedback training, for example "I believe the training helped to improve my attention", "I enjoyed the training", "The experimenter was trustworthy", etc. Furthermore, they were asked to guess the group condition they had been randomly assigned to. [...] After each session, we also assessed participants' motivation to continue participating in the training and their beliefs about being able to control their brain activity. [...] Regarding non-specific mechanisms, we were unable to find between-group differences in expectation towards the neurofeedback training and with respect to the evaluation of the training. |
| 3b | Report whether participants were provided with a strategy | we provided some example strategies that could be helpful to regulate rTPJ activity (e.g., strategies related to ToM, empathy, thinking, imagination of positive events, counting, etc.; see Supplementary Material 1). However, participants were encouraged to find their own individual successful strategy by trial and error. After each neurofeedback run, we asked participants to verbally report which strategies they used and how successful they rated this strategy (Likert scale ranging from 1 to 5). |
| 3c | Report the strategies participants used | The downregulation group used significantly more different strategies ($M = 8.66 \pm 2.47$) during the neurofeedback training compared to the upregulation group ($M = 6.26 \pm 3.24$; $t(42.11) = 2.82$, $p = 0.007$, $d = 0.87$). Figure 7 shows the distribution of strategies as reported by the participants of both groups. Fisher's exact Chi-square test revealed no significant association between the group and reported strategies ($p = 0.982$), indicating that similar strategies were used for both upregulating and downregulating TPJ activity. Table S8 shows the percentages of strategies relative to the total number of strategies reported per group and their mean success rating. In total, most strategies were reported to be more successful in the upregulation group (mean success rating: 3.35) than in the downregulation group (mean success rating: 2.74), and socio-cognitive strategies and positive mental imagery were reported most frequently in both groups (see Supplementary Material 5). |
| 3d | Report methods used for online-data processing and artifact correction | See section: Real-time fNIRS data processing (online analysis) |
| 3e | Report condition and group effects for artifacts | *Condition and group effects for artifacts were not measured, or not reported in the manuscript* |
| **Feedback specifications** | | |
| 4a | Report how the online-feature extraction was defined | See section: Real-time fNIRS data processing (online analysis) |

| | | |
|---|---|---|
| 4b | Report and justify the reinforcement schedule | Whenever participants exceeded this reward threshold, a green frame appeared around the feedback display, indicating that their regulation was earning an incentive. The total amount earned on each trial was presented on the screen at the end of the trial. This threshold was adapted according to individual regulation performance (see 2.4. for a detailed description). |
| 4c | Report the feedback modality and content | During the no-regulation condition, participants were instructed to passively look at the avatar, which maintained a neutral facial expression. During the regulation condition, real-time feedback of rTPJ activity was presented visually on a screen using a smiling avatar (social reward). Participants were instructed to regulate and make the avatar smile, which was modulated in real time by their rTPJ activation. |
| 4d | Collect and report all brain activity variable(s) and/or contrasts used for feedback, as displayed to experimental participants | See results section: 3.3 Neurofeedback regulation success |
| 4e | Report the hardware and software used | See sections: 2.3 fNIRS acquisition and 2.4 Real-time fNIRS data processing (online analysis) and subsection Statistical methods and software of section 2.5 Data processing and analysis |
| **Outcome measures - brain** | | |
| 5a | Report neurofeedback regulation success based on the feedback signal | see results section: 3.3 Neurofeedback regulation success |
| 5b | Plot within-session and between-session regulation blocks of feedback variable(s), as well as pre-to-post resting baselines or contrasts | See Figure 5, S1, S2 and S4 |
| 5c | Statistically compare the experimental condition/group to the control condition(s)/group(s) (not only each group to baseline measures) | No specific group effect or significant group × time interaction was found. |
| **Outcome measures - behaviour** | | |
| 6a | Include measures of clinical or behavioural significance, defined a priori, and describe whether they were reached | *The manuscript does not include measures of clinical or behavioural significance* |

| | | |
|---|---|---|
| 6b | Run correlational analyses between regulation success and behavioural outcomes | We found a significant negative correlation between changes in RTs in the valid trials of the reorienting of attention task and neurofeedback performance, as assessed by the number of successful runs (rho = -0.47, p = 0.045, Bonferroni corrected), indicating higher improvements of RTs in participants with more successful runs in both groups. Subgroup analysis revealed no significant effect after Bonferroni correction. For the perspective-taking task, we found a significant correlation between neurofeedback improvement (slopes) and improvements in the accuracies of NPT trials across groups (rho = -0.49, p = 0.02), indicating greater performance improvements in participants who were more successful in learning downregulation over the course of the training. This significant correlation was only observed in the downregulation group (rho = -0.71, p = 0.039, Bonferroni corrected). None of the psychosocial factors correlated significantly with behavioral outcomes after Bonferroni correction. For more details including significant correlations on the uncorrected level, see Supplementary Material 6. |
| **Data storage** | | |
| 7a | Upload all materials, analysis scripts, code, and raw data used for analyses, as well as final values, to an open access data repository, when feasible | *No additional documents related to the materials, analysis scripts, code, raw data, or final values are available for this manuscript* |
